# Dynamic representation of appetitive and aversive stimuli in nucleus accumbens shell D1- and D2-medium spiny neurons

**DOI:** 10.1101/2024.02.22.581563

**Authors:** Ana Verónica Domingues, Tawan T. A. Carvalho, Gabriela J. Martins, Raquel Correia, Bárbara Coimbra, Ricardo Gonçalves, Marcelina Wezik, Rita Gaspar, Luísa Pinto, Nuno Sousa, Rui M. Costa, Carina Soares-Cunha, Ana João Rodrigues

## Abstract

The nucleus accumbens (NAc) is a key brain region for motivated behaviors, yet how distinct neuronal populations encode appetitive or aversive stimuli remains undetermined. Using microendoscopic calcium imaging, we tracked NAc shell D1- or D2-medium spiny neurons’ (MSNs) activity during exposure to stimuli of opposing valence and associative learning. Despite drift in individual neurons’ coding, both D1- and D2-population activity was sufficient to discriminate opposing valence unconditioned stimuli, but not predictive cues. Notably, D1- and D2-MSNs were similarly co-recruited during appetitive and aversive conditioning, supporting a concurrent role in associative learning. Conversely, when contingencies changed, there was an asymmetric response in the NAc, with more pronounced changes in the activity of D2-MSNs. Optogenetic manipulation of D2-MSNs provided causal evidence of the necessity of this population in the extinction of aversive associations.

Our results reveal how NAc shell neurons encode valence, Pavlovian associations and their extinction, and unveil new mechanisms underlying motivated behaviors.

## Main

In a dynamic world, individuals are continuously flooded with sensory information of variable relevance. Our brains evolved to filter information and focus on relevant stimuli or cues predicting those stimuli. The ability to assign valence is essential to guide appropriate behavior and increase the chances of survival. Valence refers to the degree of attractiveness (positive valence) or aversiveness (negative valence) of a stimulus or outcome. From the behavioral point of view, a positive valence stimulus elicits approach whereas a negative valence stimulus triggers avoidance.

Several studies identified the NAc as essential in encoding rewarding and aversive information^1–3^. Seminal *in vivo* electrophysiological recordings report that NAc neurons innately respond to intraoral administration of appetitive unconditioned stimuli (US) such as sucrose, as well as to the aversive tastant quinine^1,2^. Interestingly, most neurons exclusively respond to positive or negative valence stimuli, though some respond to both^2^. NAc neurons also respond to reward/aversion-predictive cues (conditioned stimuli, CS)^4^, and accumbal activity is crucial for cue-outcome associations, i.e. Pavlovian conditioning^4,5^.

The NAc is mainly composed of GABAergic MSNs, segregated into those expressing dopamine receptor D1 (D1-MSNs) or D2 (D2-MSNs), though some neurons express both receptors^6^. The classical model in the field proposed a functional opposition of these two striatal subpopulations: D1-MSNs are thought to encode positive/rewarding stimuli whereas D2-MSNs encode negative/aversive stimuli^7–9^. However, this model fails to explain data from different studies^10–15^, which support a model where the two subpopulations work together to drive rewarding/aversive behaviors. For example, optical activation of either D1- or D2-MSNs supports self-stimulation, i.e. is reinforcing^12^. D1- or D2-MSN optical activation paired with a reward-predicting cue enhances motivation^16–18^. Moreover, distinct patterns of optical activation of D1- and D2-MSNs can trigger positive and negative reinforcement in the same animal^10^.

Considering the available data and the proposed role for NAc as a crucial hub connecting limbic and motor systems, one can postulate that the NAc functions as a locus where bivalent valence information is encoded and used to guide directed approach/avoidance behavior^19^. Nevertheless, despite our understanding of the contribution of this region for behavior, it has remained technically challenging to differentiate the specific role of D1- and D2-MSNs in freely behaving animals due to their similar electrophysiological properties in extracellular recordings. In this context, the development of genetically encoded calcium indicators coupled with microendoscopic 1-photon imaging enables tracking the activity of the same neurons over multiple training and stimulus’ presentation sessions. Recent studies show that NAc D1- and D2-MSNs present heterogeneous and divergent responses to rewards^20–22^. However, a recent study has proposed that NAc neurons do not signal reward or valence^23^. Therefore, decades after the recognition of the NAc as a key brain region for cue-outcome association, a critical question remains unanswered: how do neurons in this brain region encode positive and negative valence stimuli and cue-outcome associations?

To address these questions, we monitored the activity of individual NAc neurons using the cre-dependent genetically encoded calcium indicator GCaMP6f through a miniaturized microscope in D1- or A2A-cre mice (A2A was used as a marker for D2-MSNs, since cholinergic interneurons also express D2 receptor) during exposure to appetitive and aversive stimuli and during Pavlovian conditioning. Our data shows that either NAc population encode positive and negative valence unconditioned stimuli, but do not code CS valence. Surprisingly, individual neurons change their response to CS and US of either valence within trials (and across sessions), though population activity was stable and stereotypic and reliably encoded USs of opposing valence. Our data shows co-recruitment of both populations during appetitive and aversive Pavlovian conditioning, supporting a concurrent role in rewarding/aversive behaviors. We further show that D2-MSNs track US omission and extinction more prominently than D1-MSNs, and that optogenetic inhibition of D2-MSNs (but not D1-) delays extinction of aversive Pavlovian associations.

This work provides detailed functional and causal evidence regarding the role of NAc D1- and D2-MSNs in associative learning, which is of utmost importance to understand how this region contributes for motivated behaviors.

## Results

### Distinct representation of positive and negative valence stimuli in NAc D1- and D2-MSNs

To determine how NAc D1- and D2-MSNs represent stimuli of opposing valence, we recorded individual neurons in response to unsigned appetitive and aversive stimulus using a genetically encoded calcium indicator and a one-photon miniaturized microscope. For this, we injected an adeno-associated virus (AAV) expressing cre-dependent GCaMP6f in the NAc of D1-Cre or A2A-cre mice, followed by gradient index (GRIN) lens implantation in the same location (Fig. 1A-B). Six weeks later, we imaged GCaMP6f signals during a sucrose session (US1; 15μl of 20% sucrose; 39 trials) and a shock session (US2; 0.5mA, 1sec; 7 trials) (Fig. 1C). Accurate GRIN lens placement in the NAc and virus expression was assessed for all animals (Extended Data Fig. 1A).

**Figure 1.**
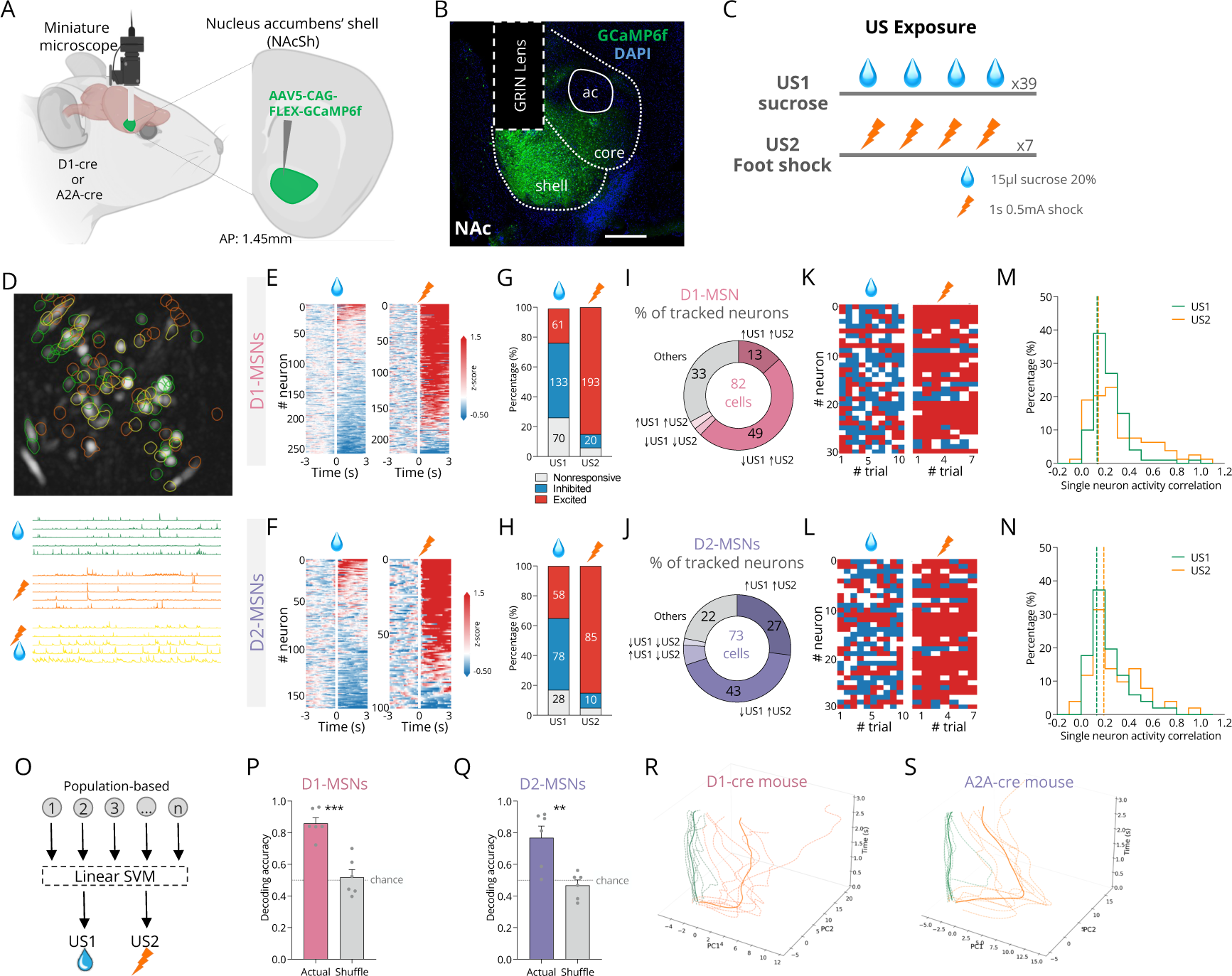
D1- and D2-MSNs encode positive and negative valence unconditioned stimuli. **(A)** Schematic representation of calcium imaging recordings of GCaMP6f in NAc D1- or D2-MSNs using 1P-miniaturized microscopes; AP: anteroposterior. **(B)** Representative expression of GCaMP6f in the NAc, depicting GRIN lens location; scale bar: 250μm. **(C)** Schematic representation of unexpected unconditioned stimulus (US) exposure sessions. US1 session consisted of 39 exposures to 15ul of 20% of sucrose with 30s ITI; US2 session consisted of 7 exposures to a 0.5mA 1s foot shock with variable ITI of 35-50s. **(D)** Top: Field of view of an example D1-cre mouse showing neurons responsive to US1 (green), to US2 (orange) and to both (yellow); bottom: single neuron traces. **(E)** Heatmaps of responses of D1-MSNs to US1 and to US2 (n_US1_=264; n_US2_=227), and of **(F)** D2-MSNs (n_US1_=164; n_US2_=100). **(G)** Percentage of excited (red), inhibited (blue) and non-responsive (gray) D1-MSNs or **(H)** D2-MSNs in response to US1 and US2. **(I)** Percentage of each type of response of neurons tracked during US1 and US2 for D1-MSNs and **(J)** D2-MSNs. **(K)** Heatmap of individual D1-MSNs or **(L)** D2-MSNs showing variable responses throughout trials. **(M)** Correlation of activity of individual D1-MSNs or **(N)** D2-MSNs in response to US1 or US2. **(O)** Cartoon depicting population-based linear support vector machine (SVM) decoder (feature: population activity during stimulus; observations: average activity during a given trial; output: US1 or US2). **(P)** Classification accuracy for every mouse using population activity of D1- or **(Q)** D2-MSNs during US1 and US2 (0-3s from US onset). The decoders were able to correctly identify sucrose and shock trials. Trajectories of trial-by-trial of **(R)** D1- or **(S)** D2-population responses to US1 and US2; Data from a representative mouse; Dots indicate stimulus onset; US1 and US2 trajectories move in orthogonal directions, supporting differential representation by D1- and D2-MSNs. Data represent mean ± SEM.

We classified a neuron as being responsive if its activity during the stimulus was significantly different from baseline (p<0.05, Permutation test). Neurons that responded to stimuli of either valence were distributed in the field of view with no obvious anatomical separation (Fig. 1D). Consistent with previous electrophysiological data^2^, sucrose consumption elicited mostly inhibitory responses in half of D1- or D2-MSNs (50% and 48% respectively); whereas shock triggered mostly excitations in both populations (85%; Fig. 1E-H).

To evaluate how the same neuron responded to stimuli of opposing valence, we analyzed the response of neurons that were tracked during sucrose and shock sessions (Fig. 1I-J) (tracked neurons activity was representative of the whole population - Extended Data Fig. 2A-D). The majority of D1- and D2-MSNs were responsive for both stimuli (67% and 78%, respectively), presenting inhibitions to sucrose and excitations to shock (Fig. 1I-J). We also characterized NAc D1- and D2-MSNs activity in response to other positive or negative valence stimuli - condensed milk and tail lift, respectively. Condensed milk led to the inhibition of 44% and 40% of D1- and D2-MSNs (Extended Data Fig. 2L-N), a smaller fraction than that observed for sucrose, which is largely consistent with the divergent response of NAc MSNs to different concentrations of sucrose^20^. Tail lift led to mostly excitatory responses in both subpopulations, in line with shock-evoked responses (Extended Data Fig. 2O-Q). These findings showing differential neuronal response to stimuli of opposing valence are in line to what is found in valence-encoding neurons from other brain regions^24,25^.

To assess the stability of neuronal responses to the same stimulus across trials, we categorized each unit into excited, inhibited or no change, based on unit average activity. Then, we calculated the fraction of persevered responses within sucrose or shock trials (analysis was performed for all recorded cells). Surprisingly, a small percentage of D1- or D2-MSNs maintained their response to sucrose in more than 70% of the trials (example neurons in Fig. 1K-L; Extended data Fig. 2E). Shock-responsive cells also changed responses throughout trials, albeit more consistent that for sucrose (Extended Data Fig. 2F). To further estimate response stability, we calculated the coefficient of correlation of activity of individual neurons across multiple trials of the same stimulus. Correlations were low for either stimulus in both MSN subpopulations (Fig. 1M-N). We also calculated Shannon’s entropy, a measure of the degree of variability in the neurons’ response. Entropy was high in sucrose trials for either D1- or D2-MSNs (Extended data Fig. 2G), whereas it was lower in shock-related activity (Extended data Fig. 2H). Importantly, the variability in individual response is not correlated with the time of the trial (Extended data Fig. 2I). Altogether, these data indicate that individual D1- or D2-MSNs do not stably encode sucrose and shock throughout time.

**Figure 2.**
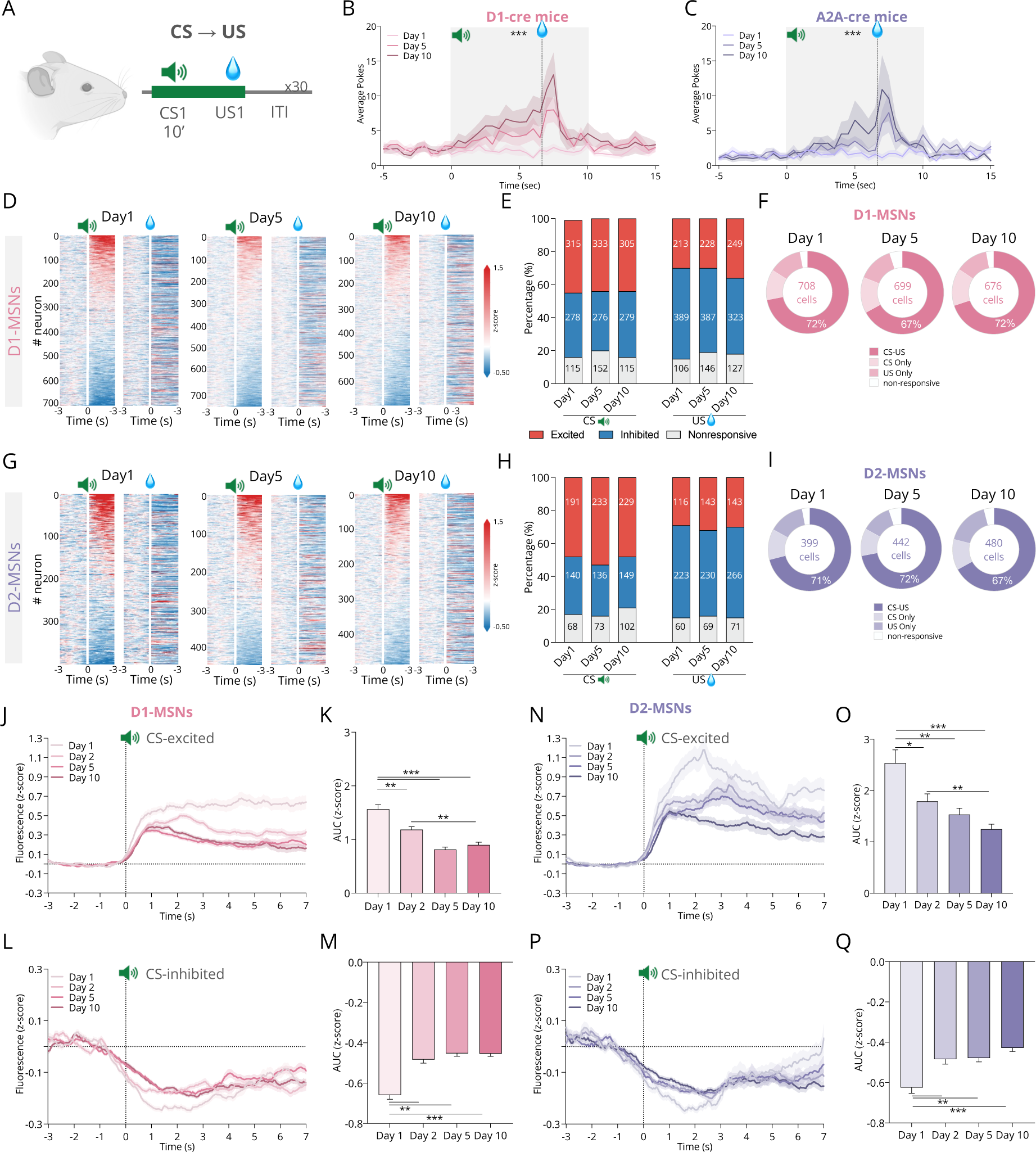
Representation of appetitive CS and US throughout Pavlovian conditioning. **(A)** Appetitive Pavlovian conditioning consisted in the association between a cue (conditioned stimulus - CS1; house light and 5-kHz tone) and the delivery of 15μl of 20% sucrose solution (US1). **(B) Average** pokes during different days of Pavlovian conditioning (day 1, day 5 and day 10) of D1-cre mice or **(C)** A2A-cre mice, showing that animals increase pokes throughout learning. **(D)** Heatmaps representing D1-MSN (n_day1_=708; n_day5_=761; n_day10_=699) or **(G)** D2-MSN (n_day1_=399; n_day5_=442; n_day10_=480) responses to CS1 (left) and US1 (right) on days 1, 5 and 10. In heatmaps, neurons are aligned by response to the CS. Percentage of responses of **(E)** D1-MSNs or **(H)** D2-MSNs to CS1 and US1 showing stability in the % of CS and US responses in both populations. **(F)** Pie chart with the % of responses throughout time for D1- or **(I)** D2-MSNs, demonstrating that the % of CS-US-responsive neurons does not change with learning. **(J)** Peri-stimulus time histogram (PSTHs) of cue-excited or **(L)** cue-inhibited D1-MSNs or **(N-P)** D2-MSNs tracked on different days of learning. **(K-M)** Area under the curve (AUCs) of mean cue response (0-3s from cue onset) for D1- or **(O-Q)** D2-MSNs over the days; there is a substantial decrease in the magnitude of response to the cue of CS-excited neurons (and less for cue-inhibited) from day 1 to day 2, suggesting that part of the cue signal is due to novelty. The magnitude of response of CS-excited D2-MSNs during the cue period is higher than in D1-MSNs. Data represent mean ± SEM.

Next, we trained a support vector machine (SVM) decoder to quantify how well individual neuron mean activity would predict trial type, i.e. distinguish sucrose from shock trials. There was a high variability in the accuracy of individual D1- or D2-MSNs (Extended Data Fig. 2J-K). Since individual NAc neurons appear to present time-dependent changes in coding properties, it is plausible that information is represented at the population level rather than at the individual level. Thus, we trained a new decoder using population responses (Fig. 1O), which accurately predicted sucrose and shock trials for D1- or D2-MSNs (86% and 77% accuracy, respectively; Fig. 1P-Q). We next sought to understand if the population represented the identity of individual stimuli or the valence of the stimuli. We observed that the decoder poorly distinguished between two positive valence stimuli - condensed milk and sucrose, though it accurately distinguished sucrose from tail lift (negative valence) (Extended Data Fig. 2R).

To further characterize how NAc activity to opposing valence stimuli unfolds over time, we calculated the neuronal trajectory, which describes the temporal evolution of neural population activity. For this, we examined trial-averaged trajectories of D1- or D2-population activity during each US. The trajectories of D1- or D2-MSNs during US1 moved in a distinct direction from US2 (Fig. 1R-S), supporting different representation of US1 and US2 by NAc neurons.

These results indicate that despite individual neuronal variability, NAc D1- and D2-population activity contains sufficient information to code the identity of positive and negative valence stimuli.

### Representation of appetitive CS and US in NAc neurons during associative learning

After determining that NAc neurons distinguished opposing valence stimuli, we then sought to characterize how these neurons would respond during associative learning, in which an initially neutral cue acquires valence and motivational value with learning. Studies performed in other brain regions involved in Pavlovian conditioning have shown that neurons undergo dynamic modifications during learning, with the development of cue responses and/or transforming existing responses^26,27^. However, it is important to refer that cue-locked neuronal responses can reflect valence attribution, but may also reflect other features such as salience or prediction error^23^.

To understand how NAc neurons’ response to CS dynamically change with learning, we imaged animals during an appetitive Pavlovian associative learning task (Fig. 2A), in which animals learn to associate an auditory and visual cue (conditioned stimulus, CS1) with the delivery of sucrose (US1). As trial exposures progressed, mice increased the number of magazine pokes and presented reduced latency to obtain the reward after delivery, indicating successful learning (Fig. 2B-C; Extended Data Fig. 3A). We were able to record several hundreds of neurons for both populations throughout the days of conditioning (Figure 2D-I). Throughout conditioning, 44% of D1-MSNs (Fig. 2D-F) and 48% of D2-MSNs presented excitations to CS1 (Fig. 2G-I). Around 1/3 of D1- or D2-MSNs presented cue inhibitions. Unexpectedly, the % of neurons that respond to CS1, US1 or both stimuli did not change significantly throughout learning (Fig. 2F, 2I; Extended Data Fig. 3B). The proportion of cue-excited or cue-inhibited neurons was remarkably stable throughout days, which contrasts with the amygdala, where an amplification in CS- (and US-) responsive neurons throughout learning was found^26,27^.

Learning can also link CS and US representations increasing correlation of responses throughout time^26^. Hence, we evaluated how CS responses were correlated with US responses on early, middle, and late learning stages (day 1, 5 and 10, respectively). Contrary to what was expected, no increase in correlation between CS and US responses for either NAc D1- or D2-MSNs was found, even when we restrict to CS-US-responsive neurons (Extended Data Fig. 3C-D).

To better understand what happens to the activity of MSNs during learning, we plotted the average activity of cue-excited and cue-inhibited neurons during different learning stages. The magnitude of response of cue-excited (but not cue-inhibited) neurons was higher in D2-MSNs in comparison to D1-MSNs (Fig. 2J, 2N), suggesting a differential contribution of these populations in cue encoding. Surprisingly, we observed a reduction in the magnitude of response of cue-excited and cue-inhibited D1- or D2-MSNs from day 1 to day 2 (Fig. 2J-Q), suggesting that part of the observed changes are due to novelty. In further support, when we compare the cue activity of the first trials with the latest trials of day 1, we observe a decrease in the magnitude of excitatory and inhibitory responses for D1-MSNs, though not significant for D2-MSNs (Extended Data Fig. 3E-F). Nevertheless, novelty *per se* does not explain the robust CS responses observed on later stages of conditioning. Recent work has proposed that NAc D2-MSNs encode valueless prediction error or signal errors^23,28^. In prediction error neurons, cue responses should develop with learning and the response to the US decrease as it becomes more predictable by the CS^29^, which we do not observe in either D1- or D2-MSNs.

Altogether, our data shows NAc cue- (and US-) evoked neuronal activity does not evolve with time, suggesting that cue responses may reflect other features of the stimulus, rather than classical prediction errors.

### Drift in the representation of appetitive stimuli in NAc neurons throughout days

Next, we took advantage of the ability to monitor individual cells throughout time to allow a more comprehensive insight on how individual neurons change throughout associative learning. For this, we followed individual neurons on days 1, 5 and 10 (272 D1-MSNs; 172 D2-MSNs) corresponding to early, mid and late training stages (representative animal in Fig. 3A; all neurons in Extended Data Fig. 2D-I, tracked neurons in Extended Data Fig. 4A-F).

**Figure 3.**
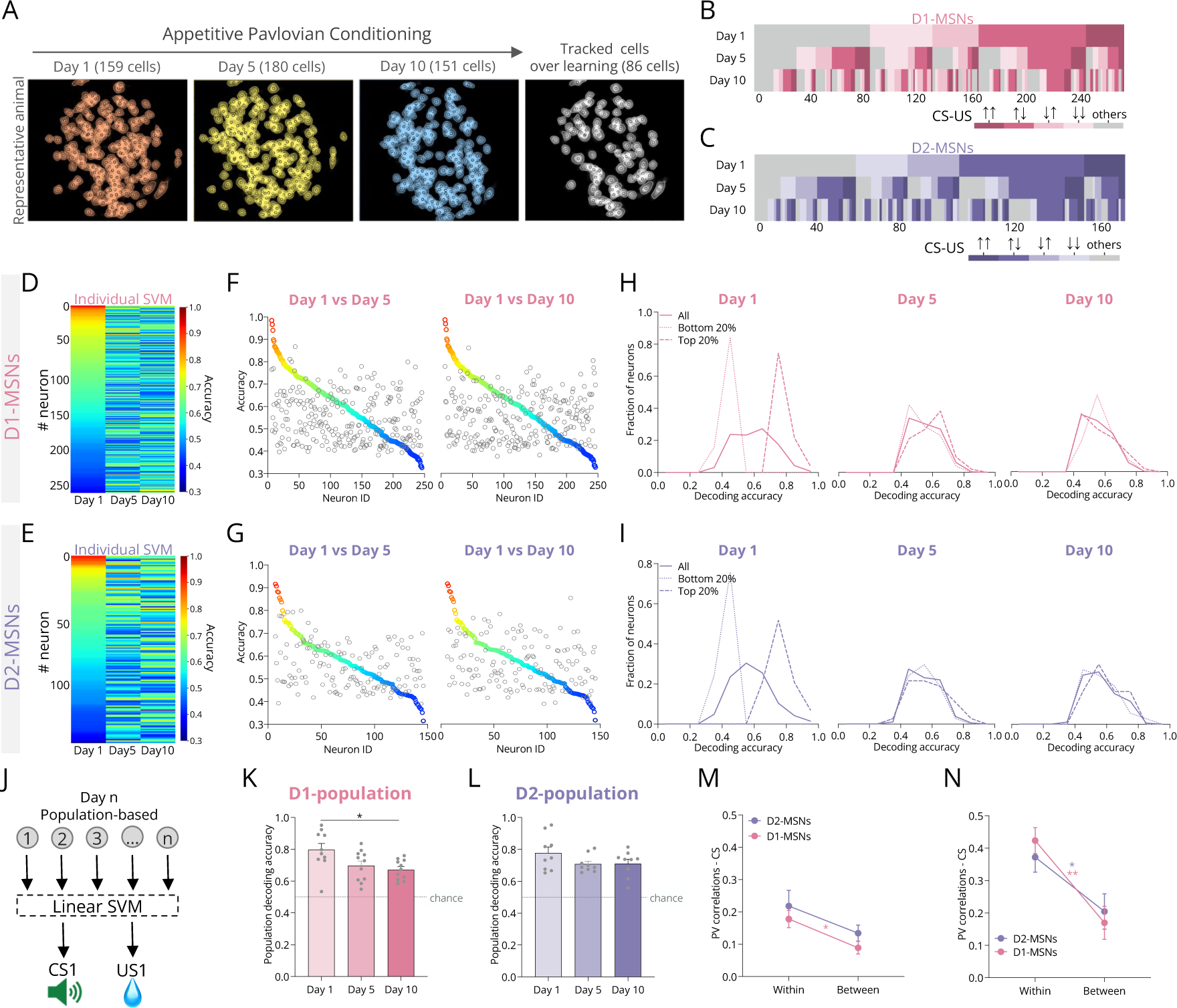
Population coding of CS1 and US1, despite individual neurons’ variability within and between sessions. **(A)** Field of view of a representative animal used for neuronal tracking during appetitive Pavlovian conditioning. **(B)** Percentage of each type of response of tracked D1-MSNs or **(C)** D2-MSNs on day 1, 5 and 10 of conditioning, showing variability in individual CS1-US1 responses over days (n_D1_= 272, n_D2_= 172). **(D)** Data from individual D1-MSNs’ activity was used to train a SVM decoder for the identification of CS1 and US1 (features: single neuron activity during stimulus; observations: average activity during given trial, output: CS1 or US1). Neurons are sorted and colored by decoding accuracy on day 1; the same was done for **(E)** D2-MSNs. **(F-G)** D1- or D2-MSNs with higher decoding accuracy on day 1 did not necessarily have relevant decoding accuracies throughout days. Colored dots represent accuracies on day 1 and grey dots represent the decoding accuracy of the same cells in days 5 or 10**. (H-I)** Decoding accuracies for all D1- or D2-MSNs and for top or bottom 20% cells in terms of accuracy. The decoding accuracy of the top 20% cells on day 1 largely overlapped with the entire population on days 5 and 10 for both populations. **(J)** Cartoon depicting Population-based SVM decoder (features: population activity during stimulus; observations: average activity during given trial, output: CS1 or US1). **(K)** SVM classification accuracy using population activity during CS1 and US1 (0-3s from CS or US onset) for D1-MSNs or **(L)** D2-MSNs. The population decoders efficiently identified CS1 and US1 events for both populations. **(M)** Population vector correlation of CS1 responses of D1-or D2-MSNs within and between sessions. **(N)** Population vector correlation of US1 responses of D1- or D2-MSNs, suggesting representational drift of US encoding between sessions.

The majority of D1- and D2-MSNs (∼70%) responded to both CS1 and US1 in all stages of learning (Extended Data Fig. 4C, F). In line with stochastic encoding of USs at a trial-by-trial level (Fig. 1), we also observed that D1- and D2-MSNs change their type of response to the CS1 trial-by-trial (Extended Data Fig. 4G), and between days (Fig. 3B-C; Extended data Fig. 4I-J). Supporting previous data, a decoder trained with the activity of each individual neuron had low accuracy in distinguishing CS1 and US1 (Extended Data Fig. 4K). Even the top 20% D1- or D2-MSNs with high decoding accuracies on day 1 did not perform well on following days (Fig. 3D-I).

**Figure 4.**
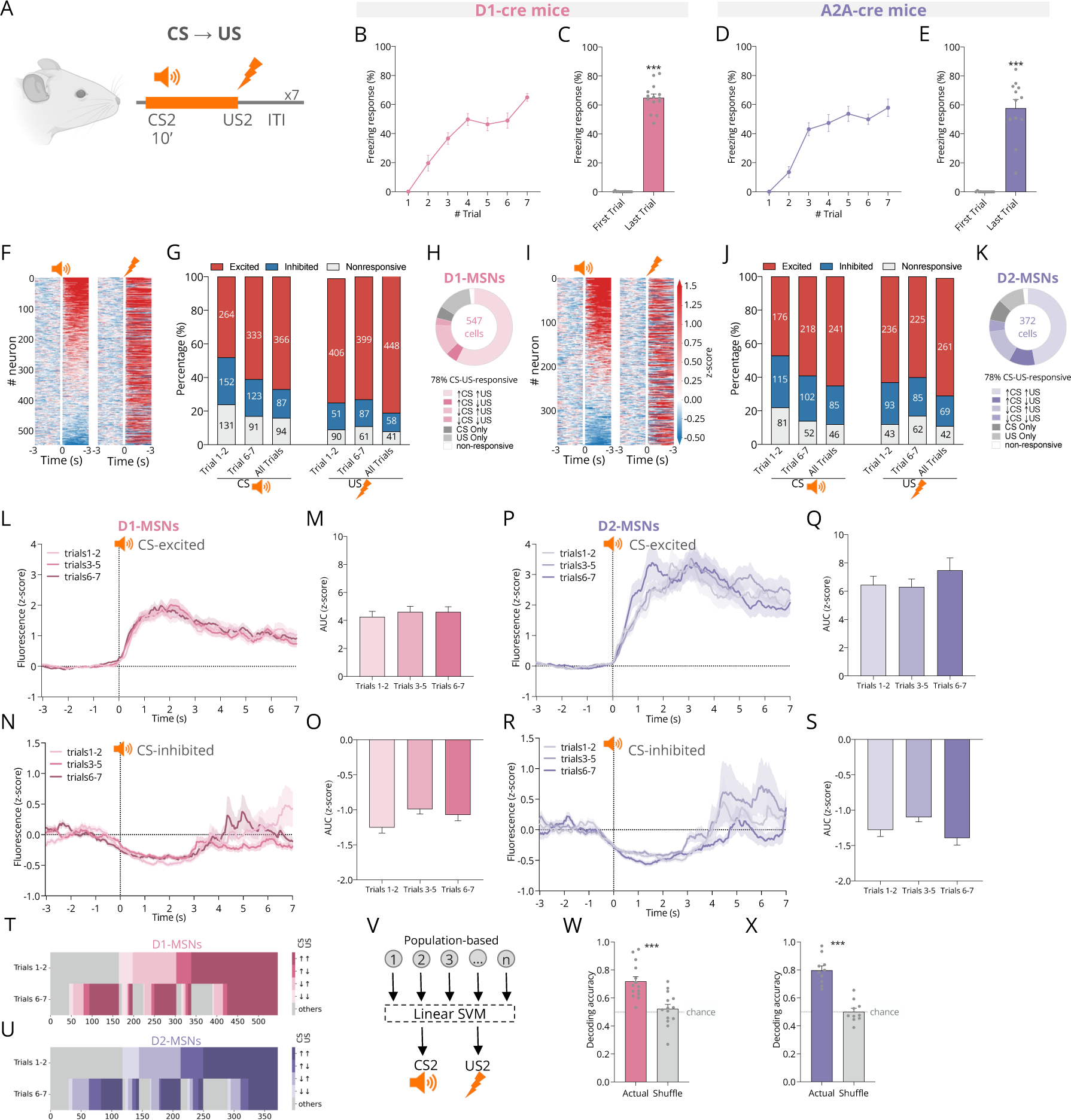
Population coding of aversive CS2 and US2 by D1- and D2-MSNs. **(A)** Aversive Pavlovian conditioning involved learning to associate an auditory cue (CS2; 3-kHz tone) with a 0.5mA 1s foot shock (US2). **(B-C)** Freezing response progression over trials of the aversive conditioning session for D1-cre and **(D-E)** A2A-cre mice. **(F)** Heatmaps representing D1-MSN responses to CS2 (left) and US2 (right). In heatmaps, neurons are aligned to response to CS2. **(G)** Percentage of each type of D1-MSNs response during the initial trials (1-2), last trials (6-7) and all trials. **(H)** Pie chart demonstrating that the % of CS-US-responsive D1-MSNs does not substantially change throughout trials. **(I)** Heatmaps representing D2-MSN responses to CS2 (left) and US2 (right). **(J)** Percentage of each type of D2-MSNs response. **(K)** Pie chart demonstrating that the % of CS-US-responsive D2-MSNs does not change throughout trials. **(L)** PSTHs of cue-excited or **(N)** cue-inhibited D1-MSNs or **(P-R)** D2-MSNs tracked on different trials. **(M-O)** AUCs of cue response (0-3s from cue onset) for D1- or **(Q-S)** D2-MSNs over the days. No major differences were observed in the magnitude of responses throughout trials. **(T)** Activity of individual D1-MSNs from the first 2 trials and to the last 2 trials, showing variability in individual responses over trials; similar findings were observed for **(U)** D2-MSNs. **(V)** Cartoon depicting population based SVM decoder (features: population activity during stimulus; observations: average activity during given trial, output: CS2 or US2). **(W)** SVM classification accuracy for every mouse using population activity during CS2 and US2 (0-3s from CS or US onset) for D1-MSNs or **(X)** D2-MSNs. D2-MSNs population activity was more accurate in the categorization of CS2 and US2 events than D1-MSNs.

The previous data strongly suggested drift in the representation of stimuli by individual neurons, thus we hypothesized that CS and US representations were encoded at the population level. Thereafter, we trained a decoder using population responses to CS1 and US1 on days 1, 5 or 10 (Fig. 3J). For either population, the decoder efficiently distinguished CS1 and US1. Surprisingly, accuracy was higher on day 1 than on subsequent days (Fig. 3K-L).

Our data showed that despite changes in coding of individual cells, population activity patterns can be used to distinguish CS and US events on different days, so we next asked if these activity patterns would be stable throughout time. To study the stability of the population representation over timescales of minutes (within session) and days (between sessions), we computed the Pearson’s correlation between pairs of trial-averaged population vectors (PV) for each stimulus within and across sessions. We observed a reduction in PV correlation between sessions for CS and for US, in comparison to within session (Fig. 3M-N).

Altogether, these data support representation of CS1 and US1 at a population-level in both D1- and D2-MSNs and suggests that these populations exhibit representational drift over sessions.

### Representation of aversive CS and US in NAc neurons during associative learning

Subsequently, we aimed to determine if NAc MSNs were involved in negative valence Pavlovian associations, thus we trained the same mice to associate another auditory and visual cue (CS2) with the delivery of a mild foot shock (US2) (Fig. 4A). Animals’ conditioned responses, measured as freezing behavior, increases throughout trials in D1-cre and A2A-cre mice (Fig. 4B-E), indicative of learning.

We observe mostly excitatory responses to CS2 or US2 in early trials (trials 1-2) and in late trials (trials 7-8) of aversive conditioning for both populations (Fig. 4F-G, 4I-J). The majority of D1- and D2-MSNs responded to both CS2 and US2 (Fig. 4H, 4K). Regardless of the neuronal population, most neurons presented excitatory responses to CS2 and US2.

To evaluate the effect of learning cue-evoked activity, we plotted the average response of cue-excited and cue-inhibited neurons on early and late trials. The response of cue-excited or cue-inhibited neurons did not significantly change throughout time (Fig. 4L-S). However, the magnitude of excitatory response to the CS was higher in D2-MSNs than in D1-MSNs (Fig. 4L, 4P).

Next, we intended to observe if the responses to CS and US were more correlated in later trials. We observed a modest effect of learning in the correlation analysis between CS and US responses in NAc D1-MSNs, but not in D2-MSNs (Extended Data Fig. 5A-B).

As observed for appetitive conditioning, individual neurons changed their response throughout trials for both D1- and D2-neurons (Fig. 4T-U), as supported by the distribution of the correlation coefficients of single neuron activity (Extended Data Fig. 5C). Because of the observed changes in individual neurons’ response, CS2 and US2 representations are also likely encoded at population level, akin to appetitive associations. Thus, we trained a decoder using the population responses to CS2 and US2 (Fig. 4V). The D1-population decoder distinguished CS2 or US2 events from shuffle data but presented lower accuracy (Fig. 4W); conversely, D2-population activity presented higher accuracy in classifying CS and US events (Fig. 4X).

In summary, we found that NAc D1- and D2-population contains sufficient information to distinguish the identities of CS2 and US2, despite individual neuronal variability.

### Similar functional clusters between D1- and D2-MSNs during Pavlovian conditioning

After determining that D1- and D2-population activity could be used to distinguish opposing valence USs, and those events from cues, we sought to identify functional ensembles containing neurons with similar patterns of activity that could better represent the impact of learning in this region. To do this, we performed principal component analysis (PCA) on neuronal responses to each CS and US on day 1, followed by K-means clustering (Extended data Fig. 6A-D). This unbiased approach identified three remarkably similar clusters on the appetitive conditioning for D1-MSNs and D2-MSNs (Extended Data Fig. 6E-F). Since we aimed to monitor the evolution of clusters’ activity throughout learning, we performed the same clustering analysis using only neurons tracked on days 1, 5 and 10. Analogous functional clusters were found for each neuronal population (Fig. 5A-B), that represented *bona fide* the activity of the whole population (Extended Data Fig. 6E-F). Importantly, clustering analysis of either population based on the activity of day 10 originated similar functional clusters (Extended Data Fig. 6G-H), suggesting constancy of the pattern of activity at a population level.

**Figure 5.**
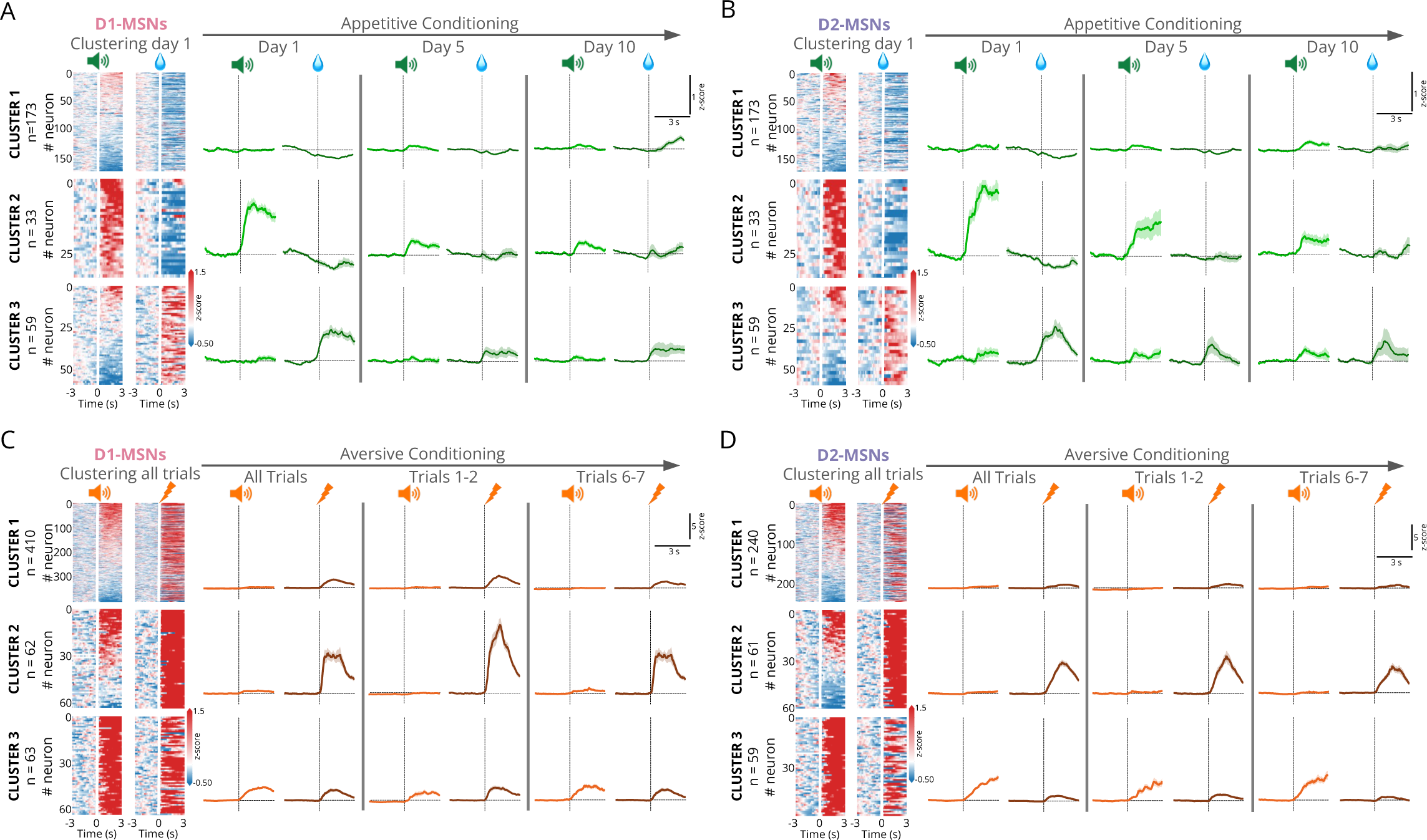
D1- and D2-MSNs form similar functional clusters during appetitive and aversive Pavlovian conditioning. Heatmaps and representation of neuronal clusters based on CS1 and US1 responses on day 1 of Pavlovian conditioning for **(A)** D1-MSNs and for **(B)** D2-MSNs, and the evolution of their response throughout days 5 and 10. Three similar functional clusters were found for each population. Neurons on heatmaps (left) are organized by CS response. **(C)** Representation of neuronal clusters based on CS2 and US2 responses during aversive Pavlovian conditioning for D1-MSNs and for **(D)** D2-MSNs, and the evolution of their response from early to late trials. Three similar functional clusters were found for each population. Neurons on heatmaps (left) are organized by CS response.

**Figure 6.**
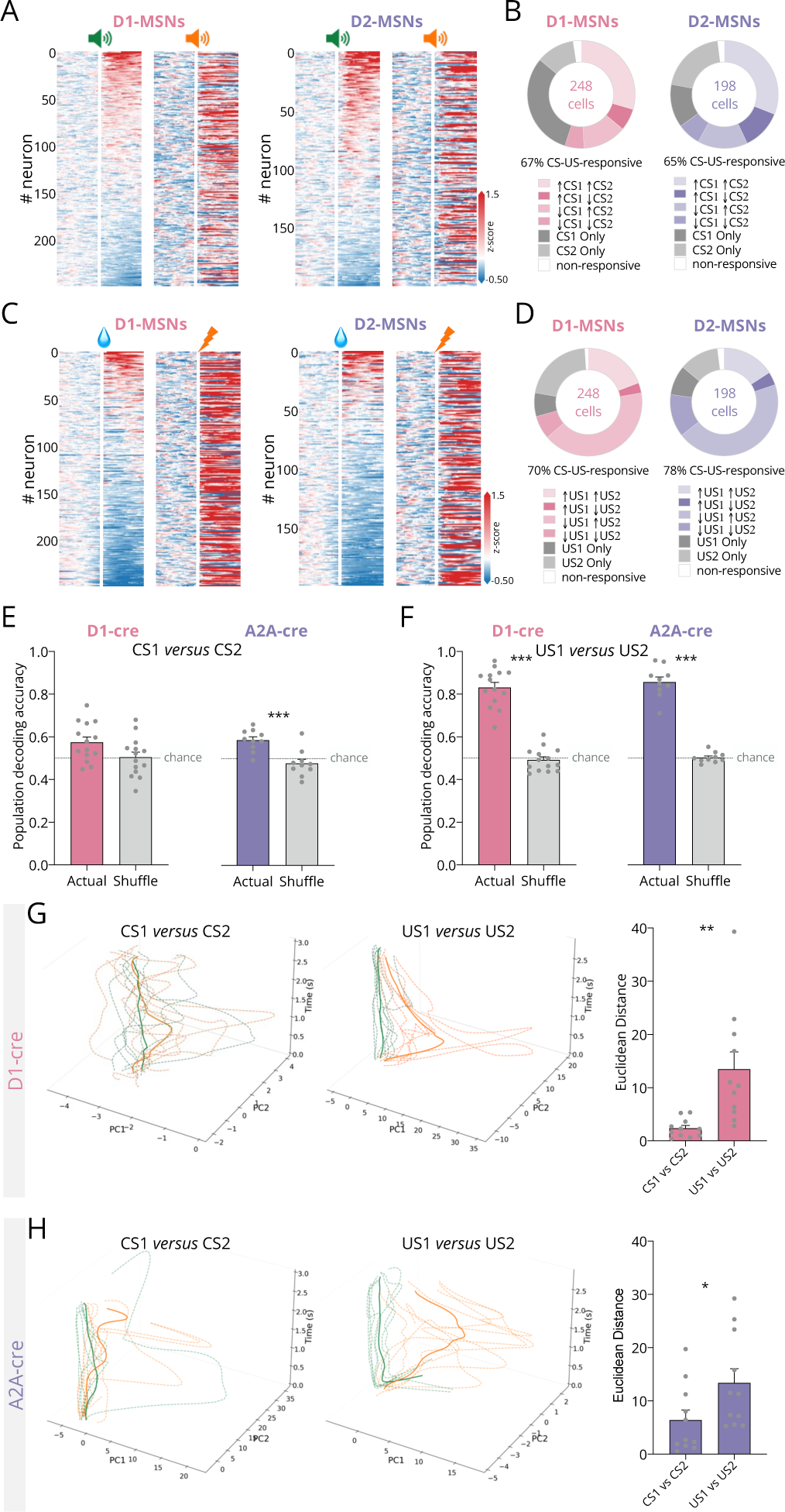
D1- and D2-MSNs encode valence of USs, but not the valence of CSs. Heatmaps representing single neuronal responses to CS1 and CS2 for D1-MSNs tracked during day 10 of appetitive conditioning and during day 1 of aversive conditioning for **(A)** D1-MSNs and **(C)** D2-MSNs. **(B)** Percentage of each type of CS1-CS2 response for both neuronal subtypes. **(D)** Percentage of each type of US1-US2 response for both neuronal subtypes. **(E-F)** The activity of D1- or D2-MSNs during CS1, CS2, US1 or US2 was used to create a population-based support vector machine decoder (features: population activity during each stimulus; observations: average activity during given trial, output: CS1 or CS2; US1 or US2). The decoder trained with D1-population data does not distinguish CSs of opposing valence, but clearly distinguishes USs of opposing valence. The decoder trained with D2-population data poorly distinguishes CS1 and CS2 trials, though it is accurate in identifying US1 and US2 trials. **(G)** Representative neuronal trajectories of one animal, depicting the trial responses to CSs or USs of opposing valence using trial-based activity data from day 10 of appetitive Pavlovian conditioning and from day 1 of aversive Pavlovian conditioning, and quantification of Euclidean distances for the trajectories of D1-MSNs. **(H)** Representative neuronal trajectories and quantification of Euclidean distances for the trajectories of D2-MSNs. In trajectories, dashed lines represent trials, full lines represent median. Note that CS1 and CS2 trajectories are intermingled, whereas USs trajectories move in distinct directions in both populations.

Then, we trained a SVM decoder using the neuronal activity of each cluster. For either population, cluster 2, which present a robust CS1 excitation and US1 inhibition, presented the higher decoding accuracy on day 1 (Extended Data Fig. 6I-J). To comprehend the temporal evolution of these clusters, we followed them throughout learning. Cluster 2 decrease the magnitude of CS response both for D1- and D2-MSNs (Extended Data Fig. 6K-L). This suggests that D1- or D2-MSNs’ cue responses do not robustly develop as a function of CS-US associative learning, as observed in VTA neurons for example^30^. Intriguingly, this cluster presents a subtle inversion in US1 response from day 1 to day 10 in both populations (Extended Data Fig. 6K-L).

We subsequently used the same unbiased clustering strategy to classify D1- or D2-MSNs into functional clusters for the aversive conditioning. For either D1- and D2-populations, three similar clusters were found (Fig. 5C-D). In the case of D1-MSNs, the cluster with higher decoding accuracy was cluster 2, that presents CS2 excitation followed by robust US2 excitation (Extended Data Fig. 6M). In the case of D2-MSNs, cluster 1 was the best performer (Extended Data Fig. 6N). Then, to observe if the activity of these clusters changes throughout conditioning, we aligned their activity from early to late trials. Throughout trials, D1-MSN cluster 2 and D2-MSN cluster 3 increase the magnitude of CS excitation, whereas US responses tend to be attenuated in later trials (Extended Data Fig. 6O-P).

Our findings indicate that D1- and D2-neuronal populations form remarkably similar functional clusters in either appetitive or aversive Pavlovian conditioning, indicating that both populations are similarly co-recruited during behavior, diverging from the classical model of striatal functional opposition.

### Valence of USs, but not of CSs, is encoded by NAc MSNs

So far, our data showed that NAc D1- and D2-MSNs can encode positive and negative valence USs (Fig. 1) and distinguish those from CSs (Fig. 3, 4). However, it was still unclear if these neuronal populations contain sufficient information to distinguish appetitive and aversive cues. To address this question, we tracked neuronal activity on day 10 of appetitive Pavlovian conditioning and on day 1 of aversive Pavlovian conditioning (Fig. 6A, 6C). More than 60% of D1- or D2-MSNs responded to both CS1 and CS2, and most of the CS-responsive neurons presented excitations, independently of CS valence (Fig. 6A-D). This observation challenges CS valence-encoding by these neurons, as one would expect differential responses to either valence cue^26,27^. Nevertheless, there was a smaller subset of neurons (24% of D1- and 27% of D2-MSNs) that responded in opposite manner for the two CSs, which could code valence. To observe if D1- or D2-neuronal activity could be used to classify event type, we used population activity patterns during CSs and USs to train a decoder. Population responses of D1-MSNs efficiently decoded sucrose and shock trials, but not the identity of CS1 and CS2 (Fig. 6E-F). Similarly, D2-neuronal population activity could be used to segregate sucrose and shock trials but poorly distinguished CS1 from CS2 (Fig. 6E-F).

We also computed the neural trajectory to CSs and USs to visualize the population activity patterns. For both D1- and D2-MSNs, CS1 and CS2 trajectories presented high trial-to-trial variability, resulting in trajectories’ overlap and low mean Euclidean distances over time (Fig. 6G-H). Conversely, US1 and US2 trajectories remained distinct and exhibited higher mean Euclidean distances, like the preconditioning phase (as depicted in Fig. 1R-S).

Our results show that the pattern of activity of D1- and D2-MSNs during CS1 and CS2 is similar, arguing against CS-valence encoding by these neurons. Since many D1- or D2-MSNs responded to both CS1 and CS2 in the same manner, this suggests that these neurons can encode CS salience, since it is expected that they respond in the same direction regardless of valence^31^.

### D1- and D2-MSNs respond to US omission, with D2-MSNs showing more pronounced responses

Our previous data suggest that CS responses encode other features of the stimuli rather than valence. The lack of CS-US changes throughout learning advocates against a *classical* prediction error encoding by NAc neurons. However, these findings do not preclude that NAc neurons are important for monitoring and updating outcomes and contribute for error signaling. To get further insight on this, we measured the activity of D1- and D2-MSNs in well-conditioned animals during unexpected US omission sessions.

After 10 days of appetitive conditioning, animals were subjected to a CS-US session in which reward was randomly omitted in 8 out of 30 trials. We evaluate neuronal activity during the first poke followed by lick event after CS1 (Extended Data Fig. 7A), to ensure that we align activity to consumption (or attempt to consume in the case of omission trials), and neurons were classified based on their average response to US consumption. No differences in CS1 responses were found between rewarded and omission trials, as expected, considering that animals do not anticipate US omission during the cue period (Extended Data Fig. 7B). D1- and D2-MSNs respond differently to reward delivery and in reward omission conditions. Sucrose-inhibited neurons no longer present inhibitions during omission in either population (Extended Data Fig. 7C-D), in further support of US-valence encoding by these neurons (Fig. 1). In fact, for both populations, neurons present an excitatory response to reward omission, with a more delayed response in D2-MSNs. Regarding sucrose-excited neurons, we observe that the two populations respond very differently, with D2-MSNs presenting a biphasic excitatory response during omission that was not observed in D1-MSNs (Extended Data Fig. 7C-D). This suggests that D2-MSNs can be important for error signaling, despite not behaving as a canonical prediction error neurons during conditioning (Fig. 2, Fig. 4).

We also evaluated the activity of D1- and D2-MSNs during unexpected shock omission. In the day after the aversive conditioning, animals were recorded in a session in which shock was randomly omitted in 4 out of 11 trials (Extended Data Fig. 7E-I). No major differences in CS2 responses were found between trials, in agreement with the randomness of the shock omission (Extended Data Fig. 7F). Regarding response to shock omission, we observe a robust increase in the number of inhibited neurons in both populations (Extended Data Fig. 7G). This can reflect outcome updating, though another parsimonious explanation is that these changes signal a positive valence outcome, due the omission of an expected noxious stimulus. Importantly, there was higher % of D2-inhibited neurons during shock-omission in comparison to D1-MSNs (Extended Data Fig. 7G).

Altogether, our results suggest that D2-MSNs signal US omission more robustly than D1-MSNs, supporting a model in where these neurons monitor and update outcomes.

### Differential contribution of NAc D1- and D2-MSNs during extinction learning

Continuous omission of USs will eventually extinguish the learned association. Extinction is a form of learning that is thought to involve new brain plasticity that encodes a “CS-no US” association, though there can also occur the degradation of the previous association^32^. We recorded D1- and D2-MSNs activity in extinction conditions (CS, no US). After appetitive Pavlovian conditioning, animals were subjected to three extinction sessions, in which the CS1 was presented but no US1 (sucrose) was given (Extended Data Fig. 8A). D1- and A2A-cre animals present reduced conditioned responses since extinction session 1 (Extended Data Fig. 8B-C). Regardless of the neuronal population, most recorded cells presented excitatory responses to CS1 during extinction days (Extended Data Fig. 8D-G). These results were also confirmed with data of neurons tracked on day 10 of Pavlovian conditioning and on extinction days 1 and 3 (Extended Data Fig. 8H-K).

Next, we evaluated the responses of D1- and D2-MSNs during extinction of aversive Pavlovian associations. After aversive Pavlovian conditioning, animals were subjected to nine days of extinction, being exposed to CS2 but with no shock (US2) delivered (Fig. 7A). Freezing was used as a measure of conditioned responses. Throughout extinction sessions, both D1- and A2A-cre mice reduced freezing behavior, as expected (Fig. 7B-C). Of notice, there was an uneven response of D1- and D2-MSNs to the CS in extinction conditions (Fig. 7D-H – tracked neurons; data from all recorded neurons in Extended Data Fig. 9A-E), since the % of excitatory and inhibitory responses throughout extinction is divergent. The percentage of CS2-excited D1-MSNs increases on extinction day 1 from 63% to 80% but substantially decreases on extinction days 5 and 9 (Fig. 7D). Conversely, the % of D1-inhibited cells increases throughout extinction. In contrast, D2-MSNs preserve the percentage of CS2-excited or CS-inhibited responses throughout extinction (Fig. 7D). When we analyzed the magnitude of response throughout extinction in tracked neurons, we observed that the CS-excitatory response was more prominent in D2-MSNs in comparison to D1-MSNs throughout days (Fig. 7E-H).

**Figure 7.**
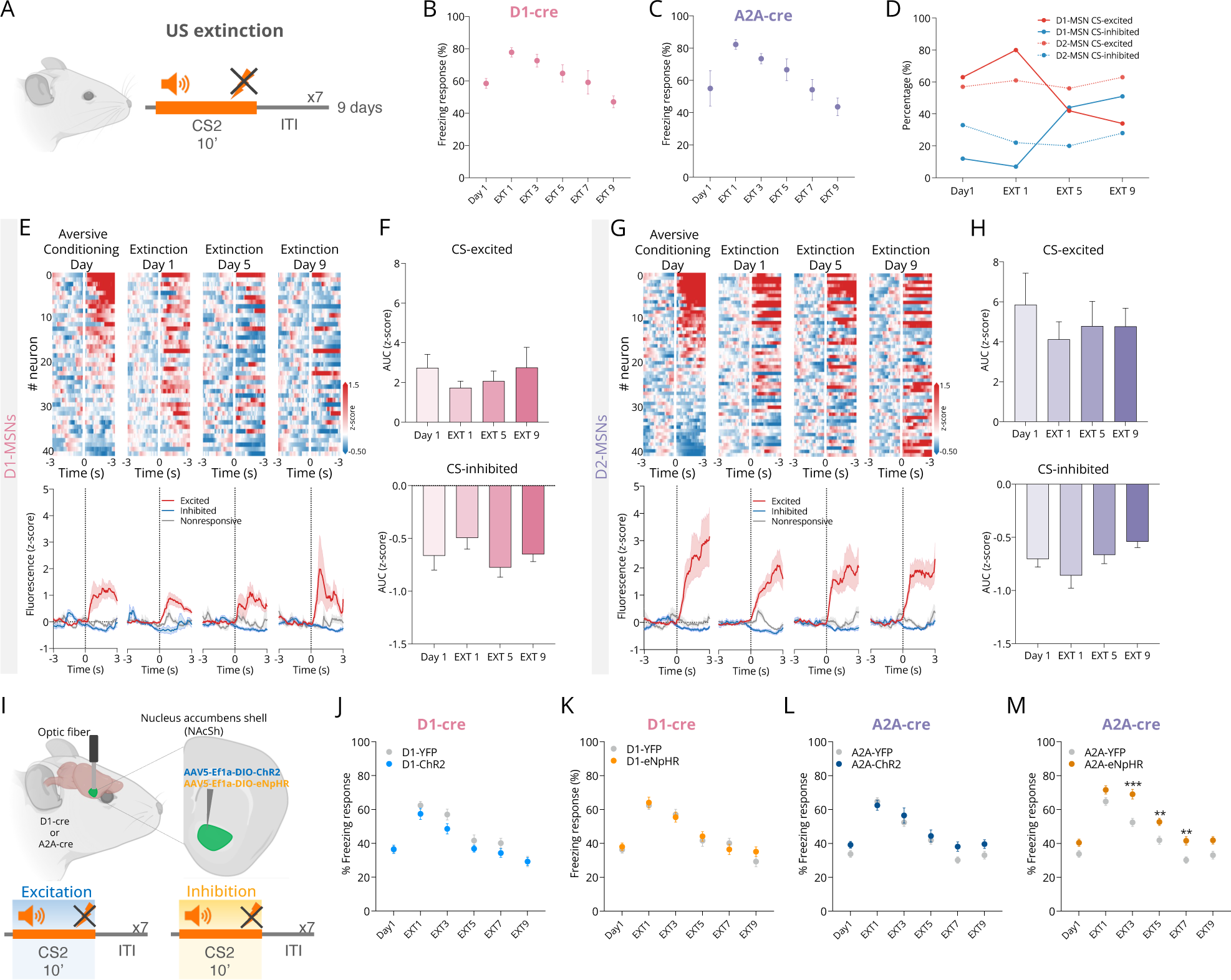
D2-MSNs are essential in the extinction of aversive Pavlovian associations. **(A)** After aversive Pavlovian conditioning, animals were exposed to 9 days of extinction (CS2, no US2). **(B-C)** Freezing responses of D1-cre or A2A-cre animals, respectively, showing a decrease in aversive conditioned responses throughout extinction. **(D)** Percentage of excitatory and inhibitory responses to CS2 in the day of aversive conditioning and in extinction sessions 1, 5 and 9 for D1-MSNs and D2-MSNs**. (E)** Heatmaps of tracked D1-MSNs or **(G)** tracked D2-MSNs showing the CS2 responses throughout days. In heatmaps, neurons are aligned to CS2 response on the conditioning day. **(F)** AUCs of D1-MSNs or (**H)** D2-MSNs CS responses throughout the days. **(I)** Scheme of the optogenetic manipulation during aversive Pavlovian conditioning. D1- and A2A-cre mice were exposed to a fear conditioning session without optogenetic manipulation. After, mice were exposed to 9 days of extinction, with optical excitation or optical inhibition of D1- or D2-MSNs during the 10 seconds of CS exposure (excitation: 25ms light pulses at 20Hz; 473nm; n_D1-YFP_=19, n_D1-ChR2_=20; n_A2A-YFP_=19, n_A2A-ChR2_=17; inhibition: constant light; 589nm; n_D1-YFP_=19 n_D1-NpHR_=17; n_A2A-YFP_=19, n_A2A-NpHR_=18). **(J)** Optogenetic excitation or **(K)** inhibition of D1-MSNs during extinction sessions does not alter the extinction of conditioned responses. **(L)** Optogenetic excitation of D2-MSNs during extinction sessions does not alter conditioned responses. **(M)** Optogenetic inhibition of D2-MSNs during CS period in extinction sessions delays the extinction of conditioned responses, as shown by the increased freezing behavior in relation to YFP animals.

Together, these results indicate that D1- and D2-MSNs have a differential response to an aversive CS under extinction conditions. D2-MSNs have a more prominent and sustained CS-excitatory response profile under extinction conditions in comparison to D1-MSNs, pointing to a more relevant role of this population in the extinction process.

### Optogenetic manipulation of NAc D2-MSNs, but not D1-MSNs, during aversive CS delays extinction of conditioned responses

The differential temporal dynamics in CS response between D1- and D2-MSNs, suggests that the two populations have distinct contributions in extinction conditions. Considering the sustained response of D2-MSNs and the magnitude of their response throughout extinction sessions, we hypothesized that by inhibiting these neurons, one could modulate the extinction association and change conditioned responses. To test this hypothesis, we injected D1- or A2A-cre animals with a cre-dependent inhibitory opsin in the NAc (eNpHR), or an excitatory opsin (ChR2) or with control YFP virus and implanted an optic fiber for optogenetic manipulation (Fig. 7I).

Animals were trained in the aversive Pavlovian conditioning, in which CS2 was followed by the delivery of a foot shock (7 pairings). After conditioning, animals were subjected to 9 days of extinction (CS2 only, no shock), in which we optically inhibited or excited D1- or D2-MSNs during the full cue period (Fig. 7I). Optical excitation or inhibition of D1-MSNs during cue period in extinction conditions did not alter the slope of extinction of conditioned responses in comparison to YFP control animals (Fig. 7J-K). Optical excitation of D2-MSNs during extinction sessions also had no effect in freezing (Fig. 7L). Remarkably, and supporting our hypothesis, optical inhibition of D2-MSNs during CS2 in extinction sessions significantly delayed the extinction of the conditioned response, since A2A-NpHR animals presented higher freezing behavior in comparison to control YFP group (Fig. 7M). Importantly, inhibition of D2-MSNs (or D1-MSNs) during the CS period of the conditioning day had no significant impact in freezing behavior (Extended Data Fig. 10A-B). No major differences in locomotor behavior were observed in any of the groups (Extended Data Fig. 10C-D).

Overall, this experiment demonstrates the essential contribution of D2-MSNs, but not of D1-MSNs, for the extinction of aversive Pavlovian associations.

## Discussion

In this study, we examined the specific features of NAc D1- and D2-neuronal responses to positive and negative valence stimuli within the same individual and decoded their role in associative learning. We show that despite stochastic encoding at individual level, NAc D1- or D2-population activity reliably encodes positive and negative valence unconditioned stimuli. The two populations form remarkably similar functional clusters during Pavlovian conditioning, supporting a model where both populations simultaneously work together to drive appropriate associative learning. However, contrary to other brain regions involved in associative learning^26,27^, cue-evoked accumbal activity does not encode valence or canonical prediction errors. We show that D2-MSNs signal reward omission and extinction more prominently than D1-MSNs, supporting a key role in monitoring and updating outcome information. In line, optogenetic inhibition of D2-but not D1-MSNs delays extinction of aversive Pavlovian associations.

Here, we demonstrate that NAc MSNs, regardless of being D1- or D2-MSNs, presented mostly inhibitions to positive valence stimuli and excitations to negative valence stimuli, in line with previous seminal electrophysiological studies of non-identified accumbal neurons in response to sucrose and quinine^2,4^. Unexpectedly, the vast majority of NAc neurons do not reliably respond to the same stimulus similarly within and between days, presenting representational drift. Still, a stable representation of USs (and CSs) emerges at a population level, despite inherent variability of individual responses. These findings are important to consider in the interpretation of studies showing that different stimuli are represented in different neurons^33^, as distinctive recruited neurons may just reflect a novel reconfiguration of the population response.

Population-level coding with single-neuron variability has also been shown for sensory representation in parietal cortex^34^ or odor coding in the piriform cortex^35^. While neuronal drift appears counterintuitive in terms of neuronal representation, it can provide the flexibility and robustness of encoding that a brain region like the NAc requires. In a constantly changing environment, the presence of multiple ways of encoding the information provides redundancy and guarantees that different dimensions of the stimulus are integrated to create a coherent representation. Drift can also allow the adjustment of the strength and structure of synaptic connections, facilitating the encoding of new information and/or refining existing representations^36^. While drift can provide these advantages, it is still necessary to reliably encode information, which we do observe at population level for D1- and D2-MSNs. One could hypothesize that in the case of the NAc, ensembles containing several neurons can be used to integrate multiple signals including valence signals arising from the amygdala^37^, sensory inputs to the NAc^38^, or context information from the hippocampus^39,40^, creating a unified and comprehensive representation of a positive or negative valence stimulus.

Lesion and pharmacological studies show that the NAc is crucial for CS-US associations and the expression of conditioned approach responses^41–44^. A key finding from our study was the remarkable similitude in D1- and D2-MSNs responses during Pavlovian conditioning, with appetitive and aversive functional neuronal clusters of each population mirroring the activity patterns of the other. These findings demonstrate that both populations act in synchrony to code rewarding/aversive information, akin to studies in the dorsal striatum showing concurrent activation of D1- and D2-MSNs in action initiation^14^. This is in agreement with behavioral studies showing that optogenetic manipulation of either D1- or D2-MSNs can drive place preference or place aversion in the same animal, depending on the pattern of activation of MSNs^45^.

In line with a relevant role for NAc in associative learning, electrophysiological studies of unidentified NAc neurons showed robust responses to CSs that develop with time^2,46,47^, consistent with a classic reward prediction error and/or valence attribution. We also found that the majority of D1- and D2-MSNs responded to appetitive and aversive CSs throughout conditioning. Interestingly, the magnitude of cue-evoked activity decreases considerably in the second day, indicating that part of the observed cue signal is due to novelty. Yet, cue-evoked activity in later stages of conditioning (and in extinction) implies that these neurons encode other features besides novelty, such as prediction errors, valence or salience^23,26,27,31,48^. A decoder trained with either D1- or D2-MSNs activity could not distinguish between opposing valence CSs, even when the association if fully established, which implies that NAc neurons encode valueless information about the cue, and that CS-valence signals are likely encoded in other regions such as the amygdala^26,27^.

The absence of temporal CS changes throughout conditioning (either by amplification of recruited neurons or increased correlation of CS-US activities) argues against a classical prediction error encoding, contrary to what has been proposed for NAc core D2-MSNs^23^. This discrepancy may be explained by anatomical specificities, considering the differential contribution of core and shell regions for Pavlovian conditioning^49–51^. Still, we observe that shell D2-neurons are activated during unexpected omission and play an important role during extinction, suggesting that they are important in error signaling/updating outcome information. One possibility is that these neurons encode salience. Salience reflects the importance of the stimulus and refers to the ability of the stimulus to capture attention and promotes associative learning^52^. It is very difficult to disambiguate salience encoding from other features, as for example unexpected US omission is an *error* and a salient event. In this framework, our data suggests that some of NAc responses are a composite of different features, so additional experiments using more sophisticated behavioral tasks are needed to isolate and track each dimension.

Despite D1- and D2-MSNs presenting remarkable similarities in activity during conditioning, their response was very distinct in extinction conditions. Sparse evidence suggests that the NAc is important for reward extinction^32,53,54^, and pharmacological blockade of dopamine receptors in the NAc impairs fear extinction learning^55^. Extinction is a fundamental form of inhibitory learning that is important for adapting to changing contingencies within the environment. The response of D2-CS-excited neurons was more robust than in D1-MSNs (both in appetitive and aversive), in line with an important role of D2-MSNs in updating information. Importantly, optogenetic inhibition of D2-MSNs (but not D1) during extinction delays the suppression of conditioned freezing response. Our results are reminiscent of another study in which optogenetic inhibition of hippocampal neurons that were active during extinction increased freezing conditioned response after extinction training^56^. These findings suggest that D2-MSNs play an important role in extinction learning, and are particularly interesting in the light of recent evidence showing that VTA dopaminergic signals to distinct NAc subregions are essential for extinction responses^53,57^.

In sum, we showed that population activity of either D1- or D2-MSNs can be used to represent and discriminate positive and negative valence stimuli. Our data strongly favors a model where the two subpopulations are co-recruited to encode CS-US associations, and elicit appetitive/aversive motivated behaviors, showing that the model of striatal functional opposition needs to be revisited. Moreover, we show that NAc neurons encode distinct features of relevant events, and that when contingencies change, D2-MSNs play a more prominent role than D1-MSNs, being essential for the extinction of aversive associations. These findings have broad implications since extinction learning constitutes a crucial component of current anxiety and post-traumatic stress disorder therapeutic interventions. Remarkably, manipulation of D2-MSNs (but not D1-) projecting to the ventral pallidum generates anxiety-like behavior^58^. Moreover, NAc dysfunction has also been found in other neuropsychiatric disorders, namely depression and addiction^59,60^, which highlights the need for further investigations to unravel the distinct contribution of different NAc neurons in the development of maladaptive behaviors.

## EXTENDED DATA – FIGURES AND TABLES

**Extended data Figure 1.**
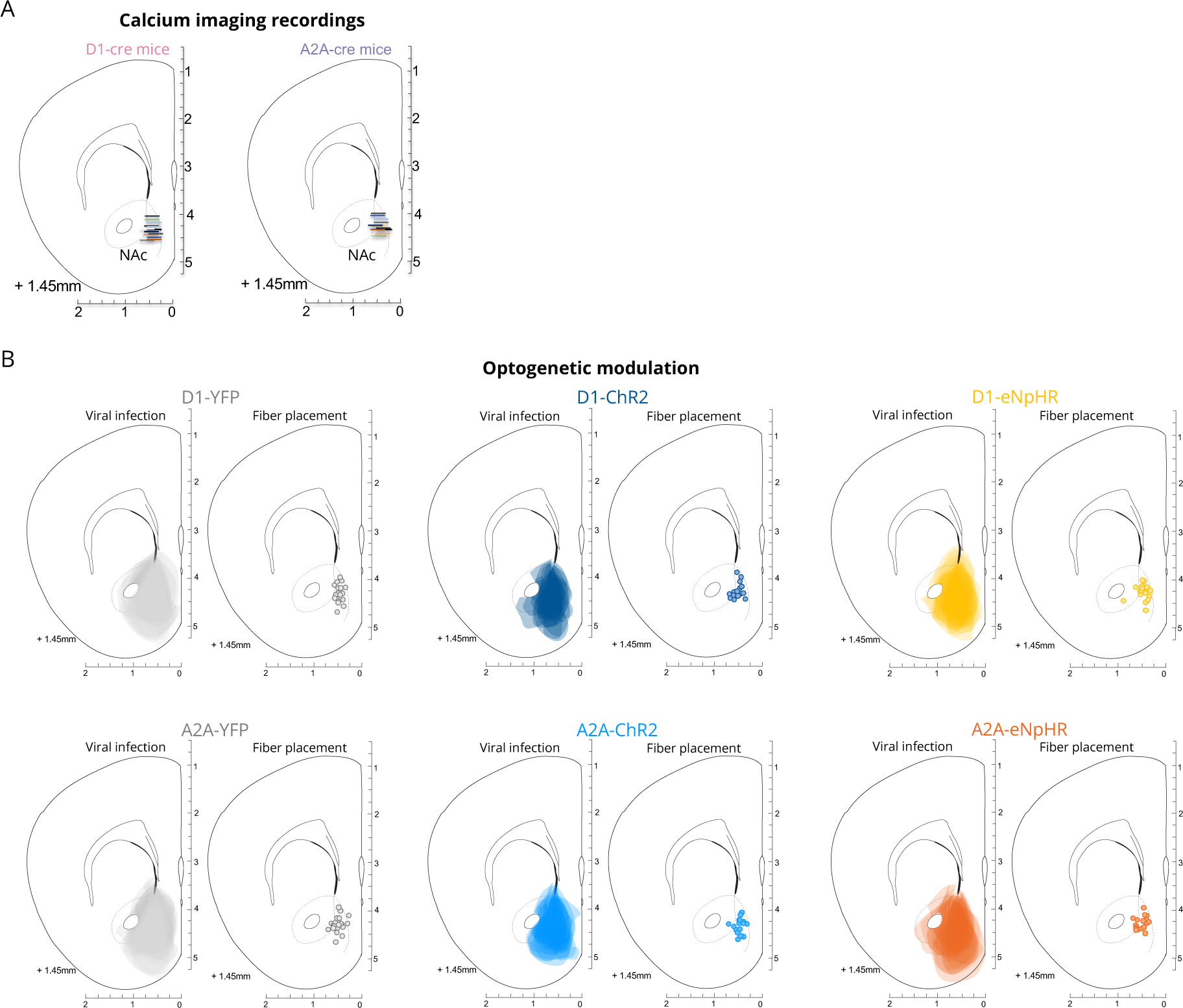
GRIN lens placements, viral expression and fiber placement for optogenetics. **(A)** Surgical placements of endoscopic GRIN lenses in mNAcSh of D1-cre animals (left) and A2A-cre animals (right). **(B)** Schematic representation of viral infection (left) and location of the tip of the fiber (right) for optogenetic manipulation.

**Extended data Figure 2.**
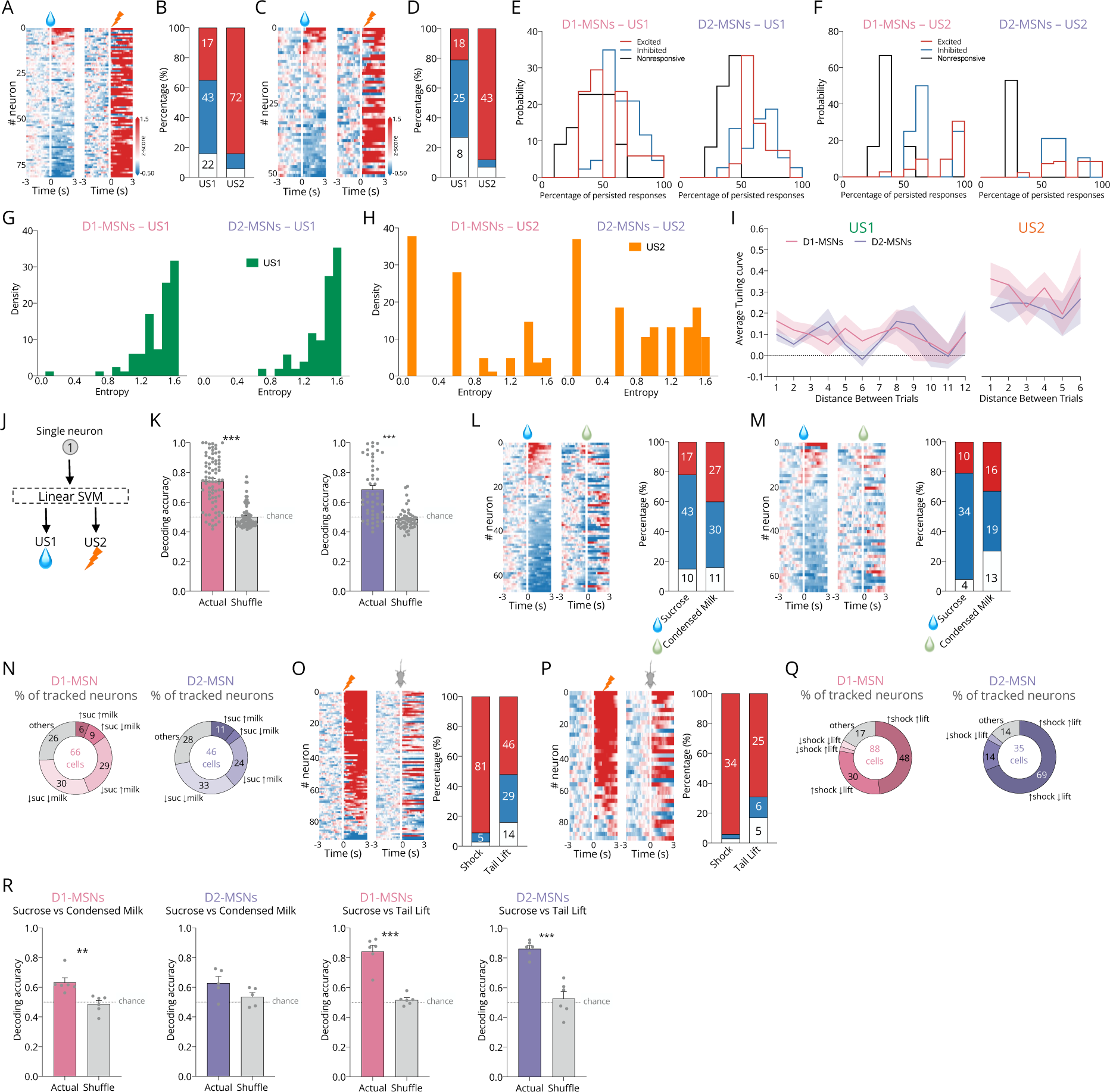
Dynamics of D1- and D2-MSN responses to positive and negative valence stimuli. Heatmaps of responses of tracked neurons to US1 (left) and to US2 (right) in **(A)** D1- and **(C)** D2-MSNs. In heatmaps, neurons are aligned in terms of their response to US1 (sucrose). **(B)** Percentage of tracked neurons that were excited (red), inhibited (blue) and non-responsive (black) for D1- or **(D)** D2-MSNs. **(E)** Percentage of neurons that present persisted responses in more than 70% of the trials to sucrose consumption. **(F)** Percentage of neurons that present persisted responses in more than 70% of the trials to shock. **(G)** Entropy during sucrose trials. **(H)** Entropy during shock trials. **(I)** Average tuning curve of D1- or D2-MSN response to US1 or US2 within session. **(J)** Data from individual D1-MSNs activity was used to train a SVM decoder for the identification of US1 and US2 (features: single neuron activity during stimulus; observations: average activity during given trial, output: US1 or US2). **(K)** SVM classification accuracy for every neuron using activity during US1 and US2 (0-3s from US onset) for D1- and D2-MSNs. **(L)** Heatmaps representing D1-MSNs or **(M)** D2-MSNs responses to sucrose and condensed milk. In heatmaps, neurons are aligned in terms of their response to sucrose. Graphs on the right represent the % of each type of response. **(N)** percentage of each type of response to sucrose and condensed milk of each neuronal type. **(O)** Heatmaps representing D1-MSNs or **(P)** D2-MSNs responses to shock and tail lift. In heatmaps, neurons are aligned to response to shock. Graphs on the right represent the % of each type of response**. (Q)** Percentage of each type of response to shock and tail lift of each neuronal type. **(R)** Population-based support vector machine decoder (feature: population activity during stimulus; observations: average activity during a given trial, output: sucrose or condensed milk; sucrose or shock). Neither population decoder distinguishes well sucrose from condensed milk trials, but accurately identified sucrose from tail lift.

**Extended Data Figure 3.**
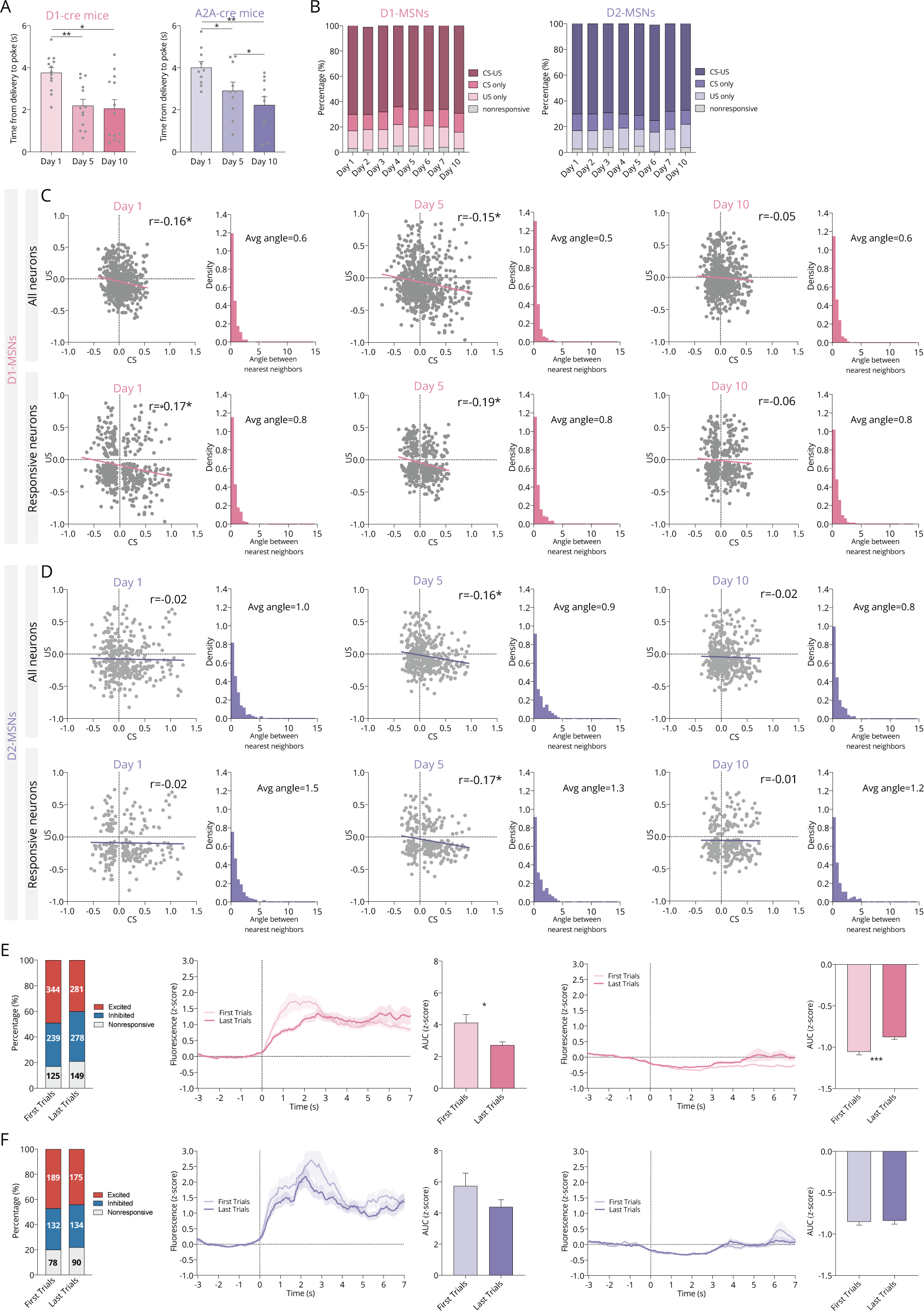
Neuronal activity of D1- and D2-MSNs over days of Pavlovian conditioning. **(A)** Throughout learning, animals decrease the time to first poke after delivery of sucrose. **(B)** Stability in the % of CS-US responsive neurons throughout days of Pavlovian conditioning. **(C)** Coefficient of correlation of activity between CS and US responses of D1-MSNs on different days of Pavlovian conditioning. Top: all neurons, bottom: considering only CS-US-responsive neurons. **(D)** Coefficient of correlation of activity between CS and US responses of D2-MSNs on different days of Pavlovian conditioning. Top: all neurons, bottom: CS-US-responsive neurons only. **(E)** Left: % of each type of CS-responses during the first 5 or last 5 trials of appetitive Pavlovian conditioning on day 1 for D1-MSNs. Middle: PSTHs showing CS-responses of excited or inhibited neurons. Right: AUCs of PSTHs. There is a decrease in the magnitude of CS-response from the first trials to the last trials. **(F)** Left: % of each type of CS-responses during the first 5 or last 5 trials of appetitive Pavlovian conditioning on day 1 for D2-MSNs. Middle: PSTHs showing CS-responses of excited or inhibited neurons. Right: AUCs of PSTHs.

**Extended Data Figure 4.**
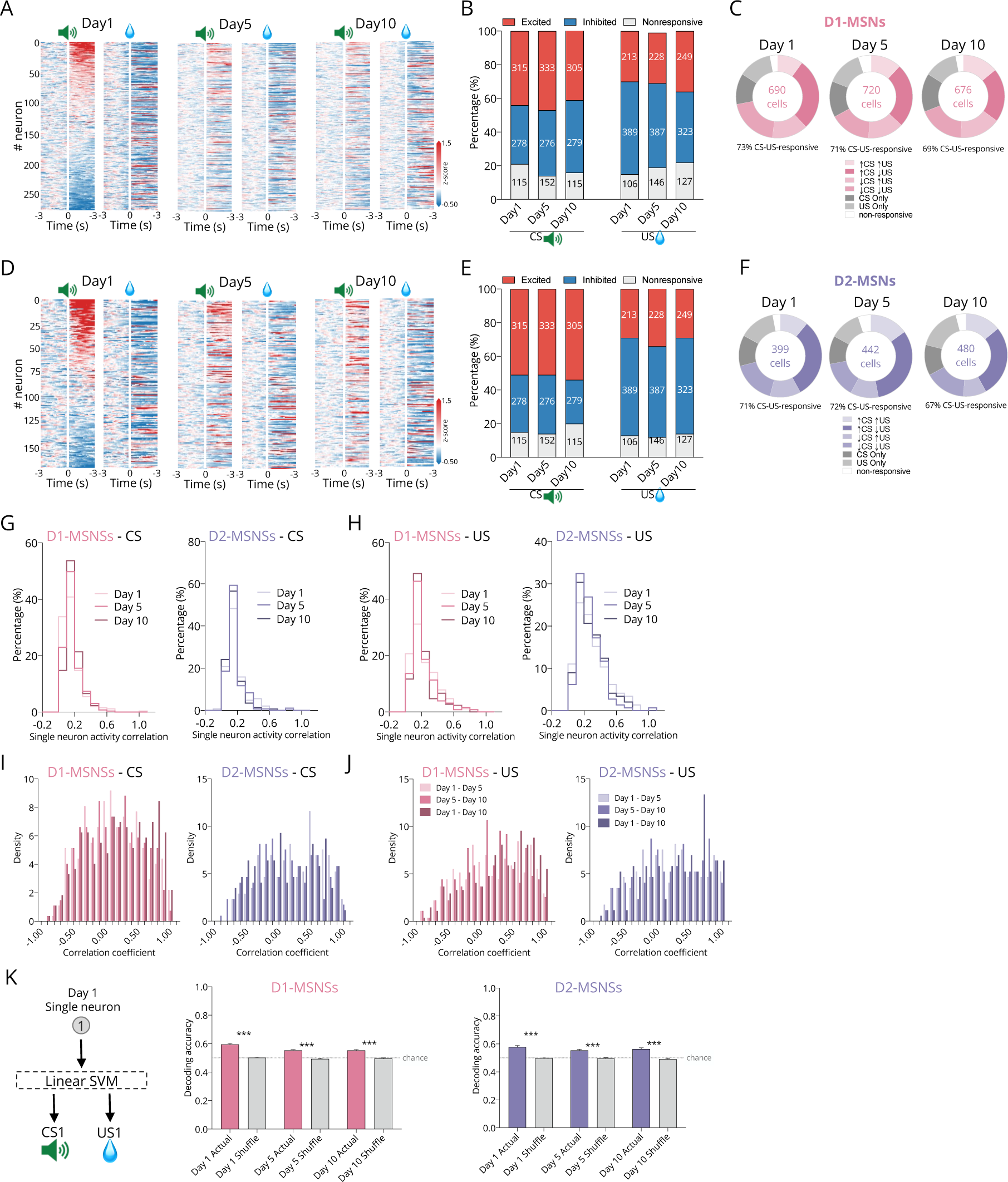
Neuronal activity of D1- and D2-MSNs tracked during Pavlovian conditioning. Heatmaps of responses of tracked neurons to CS1 (left) and to US1 (right) in **(A)** D1- and **(D)** D2-MSNs, on days 1, 5 and 10. **(B)** Percentage of tracked excited (red), inhibited (blue) and non-responsive (black) D1- and (**E)** D2-MSNs. **(C)** Percentage of each type of CS-US response in D1-MSNs or **(F)** D2-MSNs. **(G)** Single neuron CS activity correlation on day 1, 5 and 10. **(H)** Single neuron US activity correlation on day 1, 5 and 10. **(I)** Correlation coefficients of neuronal activity to CS1 between days of Pavlovian conditioning. **(J)** Correlation coefficients of neuronal activity to US1 between days of Pavlovian conditioning. **(K)** Data from individual D1-MSNs activity was used to train a SVM decoder for the identification of CS1 and US1 (features: single neuron activity during stimulus; observations: average activity during a given trial, output: CS1 or US1). SVM classification accuracy for every neuron for D1- or D2-MSNs.

**Extended data Figure 5.**
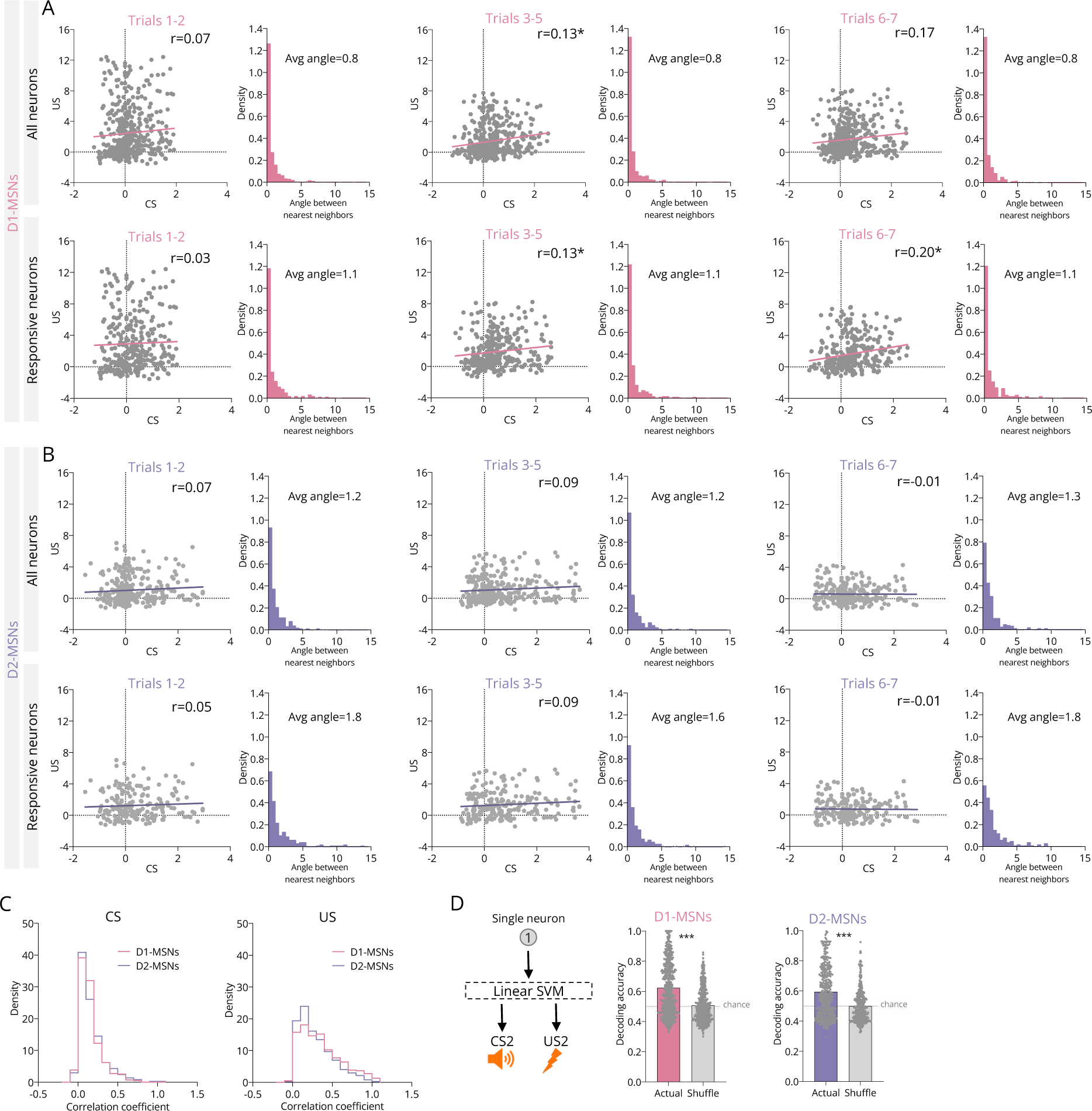
Correlations between CS and US responses during aversive Pavlovian conditioning. **(A)** Coefficient of correlation of activity between CS and US responses of D1-MSNs on early *versus* late trials in the aversive Pavlovian conditioning. Top: all neurons, bottom: considering only CS-US-responsive neurons; and for **(B)** D2-MSNs. **(C)** Single neuron CS or US activity correlation. **(C)** Data from individual D1-MSNs activity or D2-MSNs was used to train a SVM decoder for the identification of CS2 and US2 (features: single neuron activity during stimulus; observations: average activity during given trial, output: CS2 or US2). **(D)** SVM classification accuracy for every neuron for D1- or D2-MSNs.

**Extended data Figure 6.**
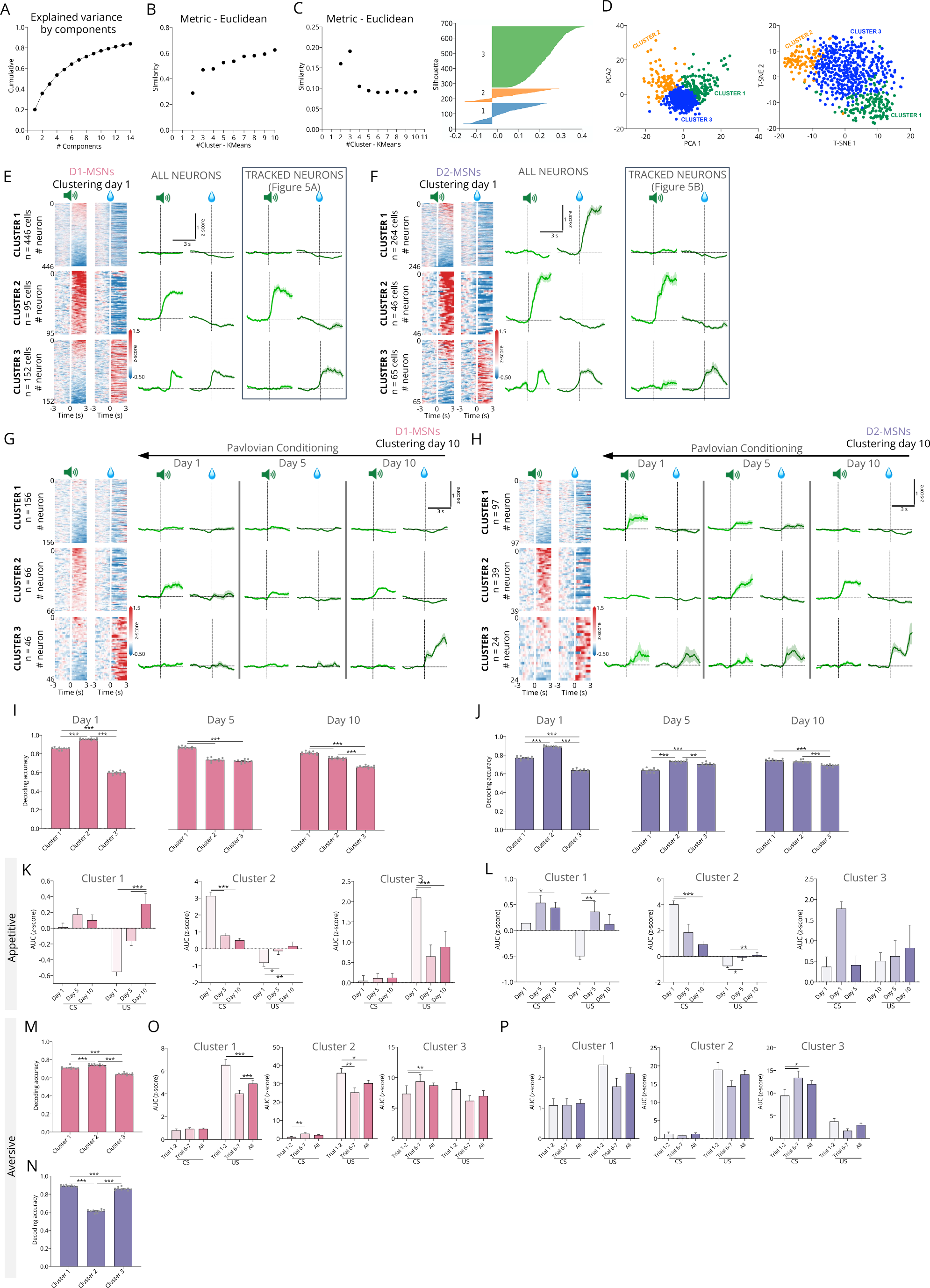
Similarity of neuronal clusters of D1- and D2-MSNs during appetitive or aversive Pavlovian conditioning. **(A-D)** Steps for clustering analysis. **A)** Plot of Cumulative variance explained by the PCA applied to CS-US average activity (z-score) over all trials. Note that, in this example, using 12 components is already sufficient to achieve at least 80% of the explained variance. **(B)** Mean cosine similarity changes with the number of clusters (n_c) chosen in the K-means clustering method; visual inspection indicates stability in similarity starting at n_c = 3. **(C)** (left) Variation of silhouette scores with the number of clusters in K-means clustering; a peak occurs at n_c = 3, aligning with the stability observed in Panel B. (right) Silhouette scores for individual clusters -red line indicates the mean of silhouette scores for n_c = 3. The combination of silhouette score analysis and cosine similarity ensure a robust determination of the optimal n_c. **(D)** (left) Clustering with n_c = 3, (upper graph); scatter plot of the first two Principal Components (PCs), where each point represents a neuron, and colors indicate the assigned cluster. (right) t-Distributed Stochastic Neighbor Embedding (t-SNE) with two components, visualizing the same data with the same color scheme. **(E)** Representation of D1 clusters formed using all neurons (left) and comparison with tracked neurons clusters’ activity (right). **(F)** Representation of D2 clusters formed using all neurons (left) and comparison with tracked neurons clusters’ activity (right). **(G)** Representation of activity of D1-clusters formed during Pavlovian conditioning based on CS1 and US1 responses on day 10 of training. **(H)** Representation of activity of D2-clusters formed during Pavlovian conditioning based on CS1 and US1 responses on day 10 of training. **(I)** The activity of D1-MSNs during CS1 and US1 of each day of each cluster was used to create a population-based support vector machine decoder (features: population activity of each cluster during the stimulus; observations: average activity during given trial, output: CS1 or US1); the same was performed for **(J)** D2-MSNs. Magnitude of CS and US responses of each cluster throughout learning, for **(K)** D1-MSNs or **(L)** D2-MSNs. For both populations, cluster 2 presents higher decoding accuracy (96% for D1-MSNs and 89% for D2-MSNs) in identifying CS and US events. **(M)** The activity of D1-MSNs during CS2 and US2 of each day was used to create a population-based support vector machine decoder (features: population activity of each cluster during the stimulus; observations: average activity during given trial, output: CS2 or US2); the same was performed for **(N)** D2-MSNs. Magnitude of CS2 and US2 responses of each cluster throughout learning for **(O)** D1-MSNs or **(P)** D2-MSNs.

**Extended data Figure 7.**
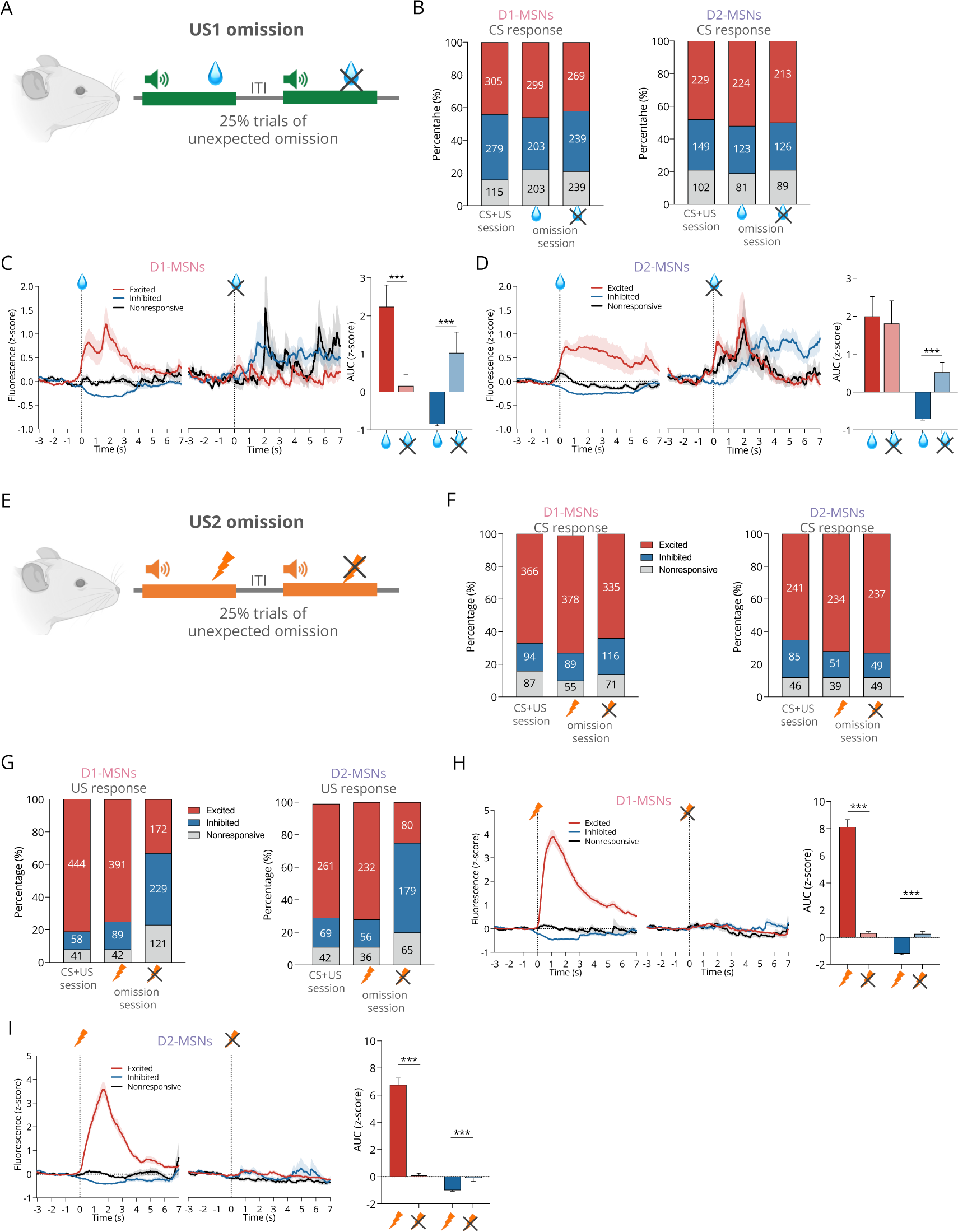
D1- and D2-MSN activity during unexpected US omission. (A) Scheme of the omission session in the appetitive conditioning. **(B)** Percentage of excited (red), inhibited (blue) and non-responsive (black) D1- and D2-MSNs to CS on appetitive Pavlovian conditioning day 10, and during reward delivery or omission. PSTHs (left) and AUCs (right) depicting **(C)** D1-MSNs or **(D)** D2-MSNs responses to reward consumption or reward omission. In PSTHs, neurons were initially classified considered their response to sucrose consumption into excited, inhibited or non-responsive and then their activity was plotted during reward consumption and in omission conditions. **(E)** Scheme of the omission session in the aversive conditioning. **(F)** Percentage of excited (red), inhibited (blue) and non-responsive (black) D1- and D2-MSNs to CS on aversive Pavlovian conditioning, and to shock or to shock omission. **(G)** Percentage of excited, inhibited and non-responsive D1- and D2-MSNs to shock and shock omission. PSTHs (left) and AUCs (right) depicting **(H)** D1-MSNs or **(I)** D2-MSNs responses to shock delivery or omission. In PSTHs, neurons were initially classified considered their response to shock into excited, inhibited or non-responsive.

**Extended data Figure 8.**
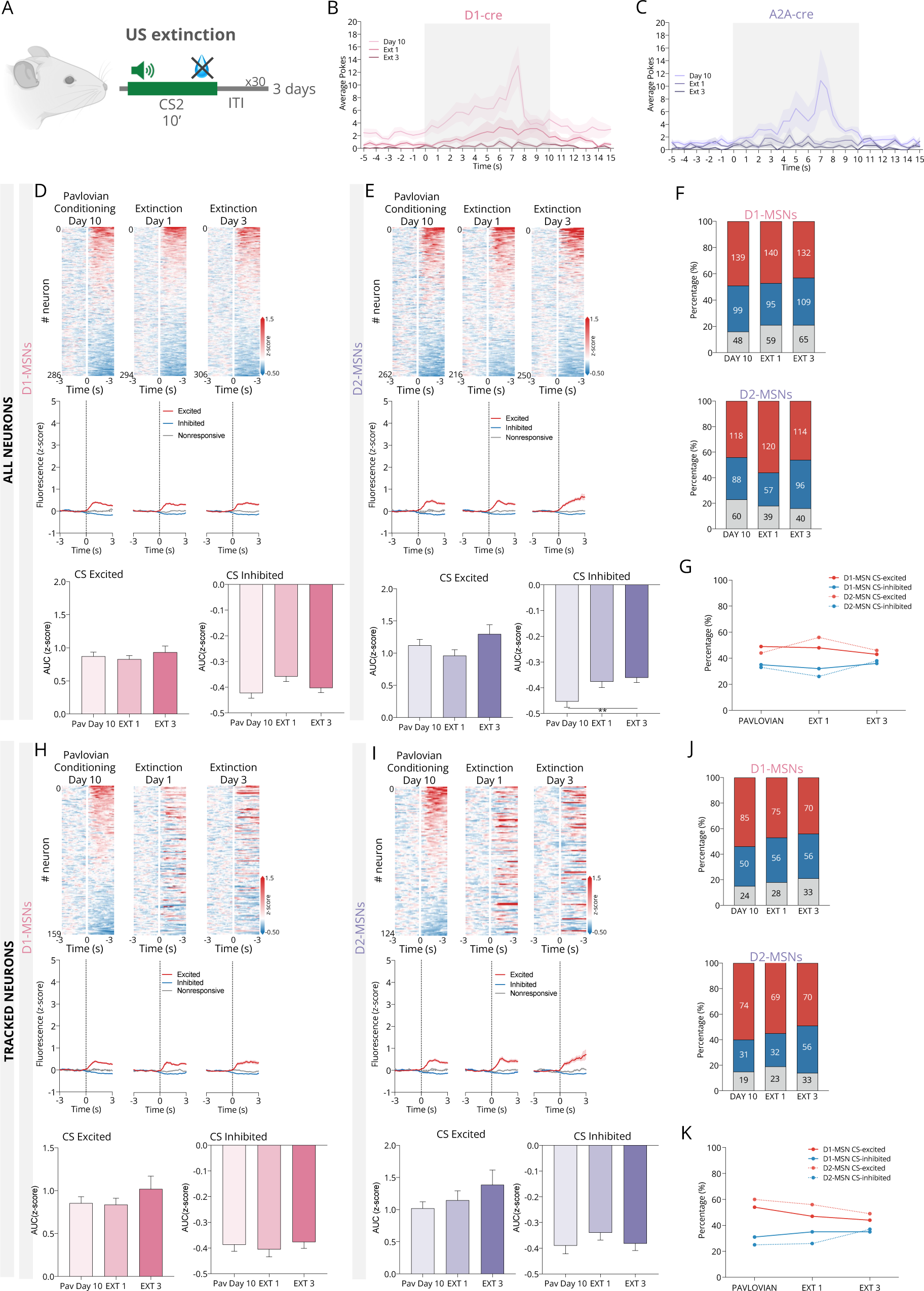
D1- and D2-MSNs activity during extinction conditions of appetitive CS-US associations. (A) Scheme of the extinction sessions of the appetitive conditioning. **(B)** Average number of pokes during Pavlovian conditioning day 10 and extinction days 1 and 3 of D1-cre mice or **(C)** A2A-cre mice, showing that animals decrease pokes throughout extinction. **(D)** Heatmaps representing all recorded D1-MSNs (n_Day10_=286; n_Day1_=294; n_Day3_=306) or **(E)** D2-MSNs (n _Day10_=262; n_Day1_=216; n_Day3_=250) responses to CS on Pavlovian conditioning day 10 and extinction days 1 and 3, and respective PSTHs (middle panel), and AUCs (bottom). In heatmaps, neurons are aligned to response to CS. **(F)** Percentage of each type of CS1 response on day 10 of appetitive conditioning and in extinction sessions 1 and 3 for D1-MSNs and D2-MSNs. **(G)** Evolution of the percentage of excitatory and inhibitory responses to CS1 in day 10 of appetitive conditioning and in extinction sessions 1 and 3. **(H-K)** The same information as depicted in D-G, but only considering neurons tracked during Pavlovian conditioning day 10 and extinction days 1 and 3 (D1-MSNs n=159; D2-MSNs n=124).

**Extended data Figure 9.**
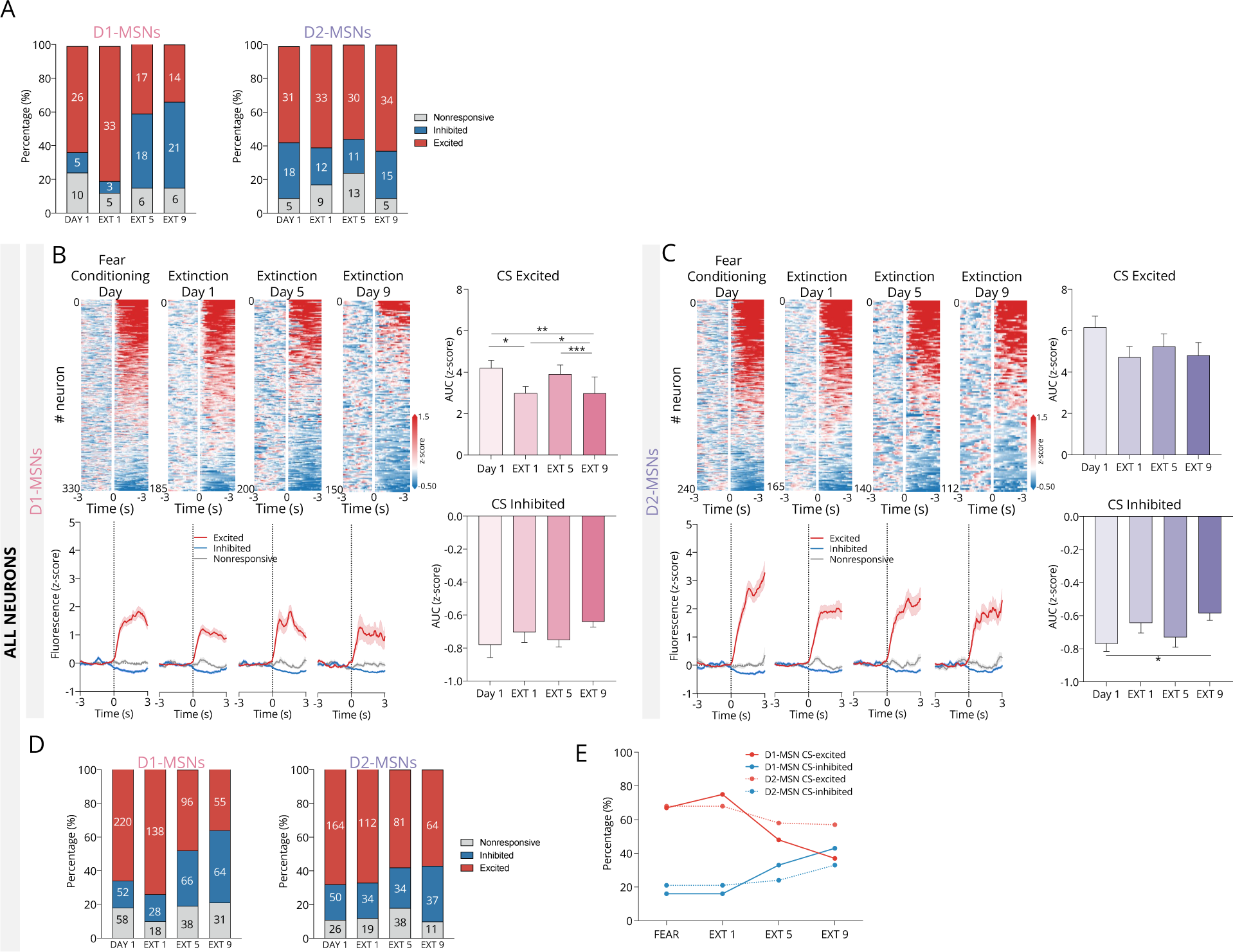
D1- and D2-MSNs activity during extinction conditions of aversive CS-US associations. (A) Percentage of CS2 response of tracked D1- and D2-MSNs on aversive Pavlovian conditioning day and during extinction days 1, 5 and 9. **(B)** Heatmaps representing all recorded D1-MSNs (n_Day1_=330; n_EXT1_=185; n_EXT5_=200; n_EXT9_=150) or **(C)** D2-MSNs (n_Day1_=240; n_EXT1_=165; n_EXT5_=140; n_EXT9_=112) responses to CS on aversive Pavlovian conditioning day and during extinction days 1, 5 and 9, and respective PSTHs (bottom) and AUCs (right). **(D)** Percentage of CS2 response of all recorded D1- and D2-MSNs on aversive Pavlovian conditioning day and during extinction days 1, 5 and 9. **(F)** Evolution of the percentage of excitatory and inhibitory responses to CS2 during extinction sessions.

**Extended data Figure 10.**
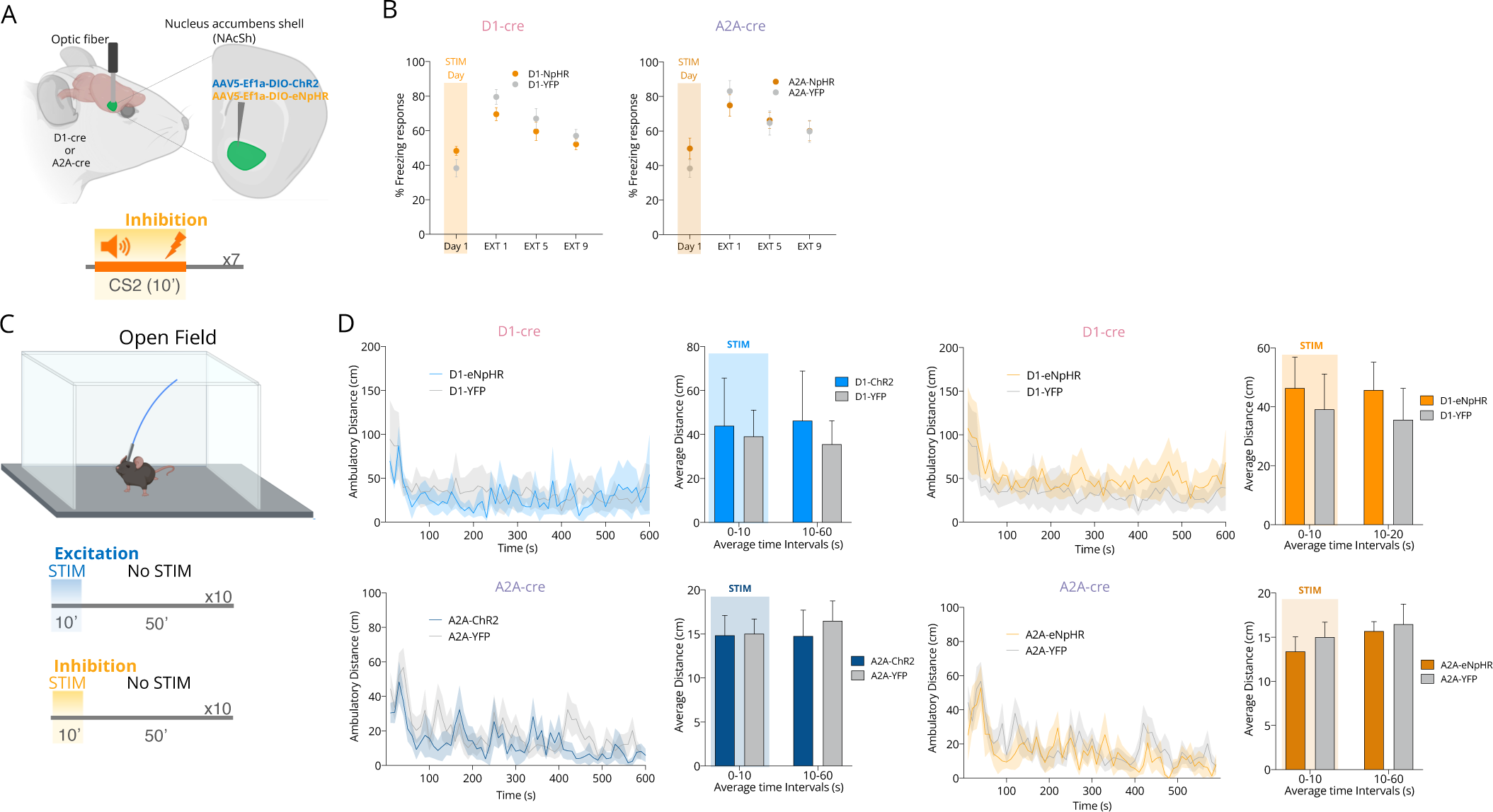
Optogenetic manipulation of D1- and D2-MSNs during conditioning and during open field. (A) Schematic representation of the experiment with optogenetic inhibition of D1- or D2-MSNs during the CS period of the conditioning day of the aversive Pavlovian conditioning. (inhibition: constant light; 589nm; n_D1-YFP_=8 n_D1-NpHR_=7; n_A2A-YFP_=6, n_A2A-NpHR_=5). **(B)** No differences in freezing behavior were observed in either group. **(C)** Schematic representation of the experiment with optogenetic inhibition during locomotion in the open field. **(D)** No major differences between average distance were found between groups. n_D1-YFP_=10 n_D1-NpHR_=8, n_D1-ChR2_=8; n_A2A-YFP_=10, n_A2A-NpHR_=8; n_A2A-ChR2_=8).

**Supplementary Table 1.** Statistical data report from main figures.

| Figure | Test | Test value | Post hoc comparison |
| --- | --- | --- | --- |
| **Figure 1** |  |  |  |
| 1P | Unpaired t test | $t_{10}=5.6$, $p=0.0002$ | |
| 1Q | Unpaired t test | $t_{10}=3.8$, $p=0.0037$ | |
| **Figure 2** |  |  |  |
| 2B | Friedman test | $F_{3,21}=34.8$, $p<0.0001$ | Dunn's<br>day 1 vs day 5, $p=0.0006$<br>day 1 vs day 10, $p<0.0001$ |
| 2C | Friedman test | $F_{3,21}=21.0$, $p<0.0001$ | Dunn's<br>day 1 vs day 5, $p=0.0036$<br>day 1 vs day 10, $p<0.0001$ |
| 2K | Kruskal-Wallis test | $KW_{4,1335}=92.3$, $p<0.0001$ | Dunn's<br>Day 1 vs. Day 2, $p=0.0091$<br>Day 1 vs. Day 5, $p<0.0001$<br>Day 1 vs. Day 10, $p<0.0001$<br>Day 2 vs. Day 5, $p<0.0001$<br>Day 2 vs. Day 10, $p=0.0012$ |
| 2M | Kruskal-Wallis test | $KW_{4,1028}=71.0$, $p<0.0001$ | Dunn's<br>Day 1 vs. Day 2, $p<0.0001$<br>Day 1 vs. Day 5, $p<0.0001$<br>Day 1 vs. Day 10, $p<0.0001$ |
| 2O | Kruskal-Wallis test | $KW_{4,889}=31.0$, $p<0.0001$ | Dunn's<br>Day 1 vs. Day 2, $p=0.0187$<br>Day 1 vs. Day 5, $p=0.0021$<br>Day 1 vs. Day 10, $p<0.0001$<br>Day 2 vs. Day 10, $p=0.0361$ |
| 2Q | Kruskal-Wallis test | $KW_{4,1335}=30.6$, $p<0.0001$ | Dunn's<br>Day 1 vs. Day 2, $p=0.0011$<br>Day 1 vs. Day 5, $p=0.0042$<br>Day 1 vs. Day 10, $p<0.0001$ |
| **Figure 3** |  |  |  |
| 3K | One-way ANOVA | $F_{2,29}=5.0$, $p=0.0140$ | Bonferroni<br>Day 1 vs. Day 10, $p=0.0163$ |
| 3M | Two-Way ANOVA | $F_{1,17}=11.7$, $p=0.0033$ | Bonferroni<br>D1-cre: within vs. between, $p=0.0033$ |
| 3N | Two-Way ANOVA | $F_{1,16}=30.2$, $p<0.001$ | Bonferroni<br>D1-cre: within vs. between, $p=0.0005$<br>A2A-cre: within vs. between, $p=0.0140$ |
| **Figure 4** |  |  |  |
| 4B | Friedman test | $F_{7,14}=57.0$, $p<0.0001$ | |
| 4C | Paired t test | $t_{12}=24.9$, $p<0.0001$ | |
| 4D | Friedman test | $F_{7,12}=47.8$, $p<0.0001$ | |
| 4E | Paired t test | $t_{11}=9.7$, $p<0.0001$ | |
| 4W | Unpaired t test | $t_{26}=4.3$, $p<0.0001$ | |
| 4X | Unpaired t test | $t_{18}=7.4$, $p<0.0001$ | |
| **Figure 6** |  |  |  |
| 6E A2A-cre | Unpaired t test | $t_{18}=4.3$, $p=0.0005$ | |
| 6F D1-cre | Unpaired t test | $t_{26}=12.2$, $p<0.0001$ | |
| 6F A2A-cre | Unpaired t test | $t_{18}=14.6$, $p<0.0001$ | |
| 6G | Unpaired t test | $t_{20}=3.4$, $p=0.0031$ | |
| 6H | Mann Whitney test | $U=28$, $p=0.0336$ | |
| **Figure 7** |  |  |  |
| 7M | Two-Way ANOVA | $F_{1,209}=57.8$, $p<0.0001$ | Bonferroni<br>A2A-eNpHR vs. A2A-YFP EXT 3, $p<0.0001$ |
|  |  |  | A2A-eNpHR vs. A2A-YFP EXT 5, p=0.0071<br>A2A-eNpHR vs. A2A-YFP EXT 7, p=0.0036<br>A2A-eNpHR vs. A2A-YFP EXT 9, p=0.0527 |

**Supplementary Table 2.** Statistical data report from extended data figures.

| Figure | Test | Test value | Post hoc comparison |
| --- | --- | --- | --- |
| **Extended Data Figure 2** |  |  |  |
| 2K D1-MSNs | Mann Whitney test | U=681, p<0.0001 |  |
| 2K D2-MSNs | Mann Whitney test | U=433, p<0.0001 |  |
| 2R D1-cre suc vs milk | Mann Whitney test | U=0, p=0.0022 |  |
| 2R D1-cre suc vs lift | Unpaired t test | t<sub>10</sub>=7.5, p<0.0001 |  |
| 2R A2A-cre suc vs lift | Unpaired t test | t<sub>10</sub>=6.7, p<0.0001 |  |
| **Extended Data Figure 3** |  |  |  |
| 3A D1-cre | Friedman test | F<sub>3,13</sub>=11.7, p=0.0029 | Dunn's<br>Day 1 vs. Day 5, p=0.0051<br>Day 1 vs. Day 10, p=0.0181 |
| 3A A2A-cre | One-Way ANOVA | F<sub>3,10</sub>=12.7, p=0.0016 | Bonferroni<br>Day 1 vs. Day 5, p=0.0500<br>Day 1 vs. Day 10, p=0.0056<br>Day 5 vs. Day 10, p=0.0397 |
| 3E AUC excited | Wilcoxon test | W<sub>281</sub>=-6317, p=0.0205 |  |
| 3E AUC inhibited | Wilcoxon test | W<sub>239</sub>=7254, p=0.0007 |  |
| **Extended Data Figure 4** |  |  |  |
| 4K D1-cre Day 1 | Mann Whitney test | U=18174, p<0.0001 |  |
| 4K D1-cre Day 5 | Mann Whitney test | U=19487, p<0.0001 |  |
| 4K D1-cre Day 10 | Mann Whitney test | U=20119, p<0.0001 |  |
| 4K A2A-cre Day 1 | Mann Whitney test | U=6251, p<0.0001 |  |
| 4K A2A-cre Day 5 | Mann Whitney test | U=6554, p<0.0001 |  |
| 4K A2A-cre Day 10 | Mann Whitney test | U=5845, p<0.0001 |  |
| **Extended Data Figure 5** |  |  |  |
| 5D D1-MSNs | Mann Whitney test | U=90526, p<0.0001 |  |
| 5D D2-MSNs | Mann Whitney test | U=45066, p<0.0001 |  |
| **Extended Data Figure 6** |  |  |  |
| 6I Day 1 | One-Way ANOVA | F<sub>2,27</sub>=1760, p<0.0001 | Cluster 1 vs. Cluster 2, p<0.0001<br>Cluster 1 vs. Cluster 3, p<0.0001<br>Cluster 2 vs. Cluster 3, p<0.0001 |
| 6I Day 5 | One-Way ANOVA | F<sub>2,27</sub>=233.5, p<0.0001 | Cluster 1 vs. Cluster 2, p<0.0001<br>Cluster 1 vs. Cluster 3, p<0.0001 |
| 6I Day 10 | One-Way ANOVA | F<sub>2,27</sub>=308.5, p<0.0001 | Cluster 1 vs. Cluster 2, p<0.0001<br>Cluster 1 vs. Cluster 3, p<0.0001<br>Cluster 2 vs. Cluster 3, p<0.0001 |
| 6J Day 1 | One-Way ANOVA | F<sub>2,27</sub>=767.7, p<0.0001 | Cluster 1 vs. Cluster 2, p<0.0001<br>Cluster 1 vs. Cluster 3, p<0.0001<br>Cluster 2 vs. Cluster 3, p<0.0001 |
| 6J Day 5 | One-Way ANOVA | $F_{2,27}=73.7, p<0.0001$ | Cluster 1 vs. Cluster 2, $p<0.0001$<br>Cluster 1 vs. Cluster 3, $p=0.0026$ |
| 6J Day 10 | One-Way ANOVA | $F_{2,27}=43.4, p<0.0001$ | Cluster 1 vs. Cluster 3, $p<0.0001$<br>Cluster 2 vs. Cluster 3, $p<0.0001$ |
| 6K Cluster 1 US | Friedman test | $F_{3,173}=37.6, p<0.0001$ | Dunn's<br>Day 1 vs. Day 5, $p<0.0001$<br>Day 1 vs. Day 10, $p<0.0001$ |
| 6K Cluster 2 CS | Friedman test | $F_{3,33}=46.8, p<0.0001$ | Dunn's<br>Day 1 vs. Day 5, $p<0.0001$<br>Day 1 vs. Day 10, $p<0.0001$ |
| 6K Cluster 2 US | Friedman test | $F_{3,33}=11.7, p=0.0029$ | Dunn's<br>Day 1 vs. Day 5, $p=0.0139$<br>Day 1 vs. Day 10, $p=0.0063$ |
| 6L Cluster 1 CS | Friedman test | $F_{3,107}=37.6, p<0.0001$ | Dunn's<br>Day 1 vs. Day 5, $p<0.0001$<br>Day 1 vs. Day 10, $p<0.0001$ |
| 6L Cluster 1 US | Friedman test | $F_{3,107}=37.6, p<0.0001$ | Dunn's<br>Day 1 vs. Day 5, $p<0.0001$<br>Day 1 vs. Day 10, $p<0.0001$ |
| 6L Cluster 2 CS | Friedman test | $F_{3,26}=32.9, p<0.0001$ | Dunn's<br>Day 1 vs. Day 5, $p=0.0002$<br>Day 1 vs. Day 10, $p<0.0001$ |
| 6L Cluster 2 US | Friedman test | $F_{3,26}=12.0, p=0.0025$ | Dunn's<br>Day 1 vs. Day 5, $p=0.0377$<br>Day 1 vs. Day 10, $p=0.0026$ |
| 6O | One-Way ANOVA | $F_{2,27}=138.8, p<0.0001$ | Cluster 1 vs. Cluster 2, $p=0.0002$<br>Cluster 1 vs. Cluster 3, $p<0.0001$<br>Cluster 2 vs. Cluster 3, $p<0.0001$ |
| 6P | One-Way ANOVA | $F_{2,27}=1067, p<0.0001$ | Cluster 1 vs. Cluster 2, $p<0.0001$<br>Cluster 1 vs. Cluster 3, $p=0.0002$<br>Cluster 2 vs. Cluster 3, $p<0.0001$ |
| 6Q Cluster 1 US | Friedman test | $F_{3,410}=23.3, p<0.0001$ | Dunn's<br>Trial 1-2 vs. Trial 6-7, $p<0.0001$<br>Trial 6-7 vs. All, $p=0.0004$ |
| 6Q Cluster 2 CS | Friedman test | $F_{3,62}=11.8, p=0.0027$ | Dunn's<br>Trial 1-2 vs. Trial 6-7, $p=0.0050$<br>Trial 6-7 vs. All, $p=0.0161$ |
| 6Q Cluster 2 US | Friedman test | $F_{3,62}=13.8, p=0.0027$ | Dunn's<br>Trial 1-2 vs. Trial 6-7, $p=0.0007$ |
| 6Q Cluster 3 CS | Friedman test | $F_{3,63}=13.8, p=0.0027$ | Dunn's<br>Trial 1-2 vs. Trial 6-7, $p=0.0055$<br>Trial 1-2 vs. All, $p=0.0029$ |
| 6R Cluster 3 CS | Friedman test | $F_{3,59}=6.4, p=0.0406$ | Dunn's<br>Trial 1-2 vs. Trial 6-7, $p=0.0388$ |
| **Extended Data Figure 7** |  |  |  |
| 7C excited | Wilcoxon test | $W_{41}=-655, p<0.0001$ | |
| 7C inhibited | Wilcoxon test | $W_{87}=3352, p<0.0001$ | |
| 7D inhibited | Wilcoxon test | $W_{138}=5145, p<0.0001$ | |
| 7H excited | Wilcoxon test | $W_{391}=-70288, p<0.0001$ | |
| 7H inhibited | Wilcoxon test | $W_{89}=3061, p<0.0001$ | |
| 7I excited | Wilcoxon test | $W_{232}=-25564, p<0.0001$ | |
| 7I inhibited | Wilcoxon test | $W_{56}=794, p=0.0009$ | |
| **Extended Data Figure 8** |  |  |  |
| 8B | One-Way ANOVA | $F_{3,21}=76.8, p<0.0001$ | |
| 8C | One-Way ANOVA | $F_{3,21}=37.0, p<0.0001$ | |
| 8E AUC inhibited | Kruskal-Wallis test | KW<sub>3,241</sub>=11.9, p=0.0026 | Day 10 vs. EXT 3, p=0.0024 |
| **Extended Data Figure 9** |  |  |  |
| 9B AUC excited | Kruskal-Wallis test | KW<sub>4,509</sub>=22.5, p<0.0001 | Day 1 vs. EXT 1, p=0.0258<br>Day 1 vs. EXT 9, p=0.0013<br>EXT 1 vs. EXT 5, p=0.0206<br>EXT 5 vs. EXT 9, p=0.0010 |
| 9C AUC inhibited | Kruskal-Wallis test | KW<sub>4,155</sub>=8.9, p=0.0309 | Day 1 vs. EXT 9, p=0.0471 |

## Online Methods

### Animals

Male and female heterozygous D1-cre (line EY262, Gensat.org) and A2A-cre (line GK139, Gensat.org) transgenic mouse lines (2-3 months of age) were used. All animals were maintained under standard laboratory conditions: an artificial 12h light/dark cycle with lights on from 8am to 8pm; with an ambient temperature of 21±1°C and a relative humidity of 50-60%. Mice were housed in type 2L home cages with a maximum of 6 mice per cage, with food (standard diet 4RF21, Mucedola, Italy) and water *ad libitum*, unless stated otherwise. After surgery, animals were maintained in pairs, without physical access to one another (using a cage divider) to avoid damaging of the implants.

Behavioral experiments were performed during the light period of the light/dark cycle. Handling was performed for 10 minutes a day, starting at least one week before behavioral experiments. Animals were habituated to behavioral apparatuses for 3 consecutive days for 15 minutes before the behavioral tasks. Sample size used in behavioral tests was chosen according to previous studies; the investigator was not blind to the group allocation during behavioral performance, but it was blind in data analysis. All procedures involving mice were performed according to the guidelines for the welfare of laboratory mice as described in the European Union Directive 2010/63/EU. All protocols were approved by the Ethics Committee of the Life and Health Sciences Research Institute (ICVS) and by the national authority for animal experimentation, Direção-Geral de Alimentação e Veterinária (DGAV; approval reference #8332). Health monitoring was carried out according to FELASA guidelines and all experimenters and animal facilities are accredited by DGAV.

### Surgeries

Surgeries were performed under sterile conditions and sevoflurane (2%–3%, plus oxygen at 1-1.5 l/min) anesthesia on a stereotactic frame (David Kopf Instruments, Model 940). Throughout each surgery, mouse body temperature was maintained at 36°C using an animal temperature controller (ATC2000, World Precision Instruments) and afterward, each mouse was allowed to recover from the anesthesia in its homecage under a heating lamp. The mouse head was shaved, cleaned with 70% alcohol and a small incision from anterior to posterior was made on the skin to allow for aligning the head and drilling the hole for the injection site.

#### Imaging

Each animal was unilaterally injected with 400 nl of AAV5.CAG.Flex.GCaMP6f.WPRE.SV40 (Addgene) into the right NAc (AP: 1.45 mm, ML: 0.6 mm, DV: 4.5 mm) using a Nanojet III Injector (Drummond Scientific, USA) at a rate of 1 nl per second. The injection pipette was left in place for 10 min post-injection before it was removed. After the injection, a 0.6-mm-diameter gradient index (GRIN) lens with a baseplate attached (Inscopix) was slowly lowered into the right mouse NAc (0.2mm per minute) directly above the injection site after a slow pre-track was made with a 26-gauge blunt needle (until 0.4mm above the target DV coordinate). Once in place, the craniotomy was closed with a low toxicity silicone adhesive (kwik-sil) and the lens was secured to the skull using dental cement (Superbond C&B kit).

#### Optogenetics

Each animal was unilaterally injected with 500 nl of cre-inducible AAV5/EF1a-DIO-hChR2(H134R)-eYFP, AAV5/EF1a-DIO-eNpHR-eYFP, or AAV5/EF1a-DIO-eYFP (UNC vector core) into the right NAc (AP: 1.45 mm, ML: 0.6 mm, DV: 4.5 mm) using a Nanojet III Injector (Drummond Scientific, USA) at a rate of 1 nl per second. The injection pipette was left in place for 10 min post-injection before it was removed. After the injection, a 0.2-mm-diameter optic fiber (Thorlabs) was slowly lowered into the right mouse NAc directly above the injection site (until 0.4mm above the target DV coordinate). Once in place, the fiber was secured to the skull using dental cement (Superbond C&B kit). At the end of the surgical procedure, mice were removed from the stereotaxic frame and postoperative care was carried out by administering analgesia (0.05mg kg-1 buprenorphine) 6h post-procedure, as well as once every 24h during three successive days. Animals were let to recover for 6 weeks before imaging recordings.

### Behavioral experiments

#### Behavioral Apparatus

Behavioral sessions were performed in a custom-made operant chamber using pyControl software and hardware (17.8cm length x 19cm width x 23cm height) within a sound-attenuating box. For appetitive stimulus (sucrose), the chamber was composed by a central magazine, to provide access to 151l of sucrose solution (20% wt/vol in water) delivered by a solenoid (for liquid dispenser), a cue-sound (70dB 5kHz), a house-light (100mA, 2.8W) installed on the top, and metallic floor. For the aversive stimulus (shock), the chamber contained a house-light (100mA, 2.8W) installed on the top of the chamber, a cue-sound (80dB 2kHz) and a cue-light installed in one side wall and a gridded floor with shocker. A computer was used to control the equipment and record the data, and a webcam (CMOS OV2710, ELP, Shenzhen, China) was used to acquire video.

### Exposure to distinct USs

#### Appetitive USs

##### Sucrose (US1)

After 3 days of habituation to the behavioral box and the miniscope, mice (n_D1-cre_=15, n_A2A-cre_=12) were exposed to 1 session of sucrose consumption, in which 151l of a 20% sucrose solution were delivered every 30 seconds, for 30 minutes.

##### Foot Shock (US2)

The same mice were familiarized to the aversive chamber apparatus for 3 days for 10 minutes with the patch cable connected. The foot shock session consisted of the unpredictable delivery of 10 mild foot shocks (0.5mA, 1 sec), separated by a random ITI (35-50 seconds).

##### Condensed milk (US3)

After sucrose session, mice (n_D1-cre_=7, n_A2A-cre_=6) were exposed to 1 session of condensed milk consumption, in which 151l of a condensed milk solution (10% sugar) were delivered every 30 seconds, for 30 minutes.

##### Tail Pinch (US4)

Mice were placed in an open arena covered with corn cob bedding and were allowed to explore the arena for 10 minutes. After that, a manual pinch to the tail was applied with an interval of 30 seconds for 5 times.

### Appetitive Pavlovian conditioning

All animals performed first the appetitive Pavlovian conditioning (protocol adapted from^56^) and posteriorly the aversive Pavlovian conditioning. After sucrose consumption session, mice started the appetitive Pavlovian conditioning in which a conditioned stimulus (CS1) consisting of a 70dB 5kHz tone and a house-light (100mA, 2.8W) was turned on for 10 seconds; 151l of 20% sucrose solution (unconditioned stimulus, US1) was made available at the 7th second after CS onset. CS-US pairings were repeated 30 times per session, with a variable inter-trial interval (ITI) of 15-35 seconds (randomly assigned). Mice underwent a total of 10 sessions of appetitive Pavlovian conditioning. The behavior apparatus and the sucrose receptacle were disinfected with 10% ethanol between animals to remove any odor. For all sessions, nose poke and licks (for half of the animals, as in one set the animals the lickometer was not properly working) data and imaging recordings were simultaneously obtained and synchronized through pyControl and IDAS (Inscopix) systems. To quantify CS-triggered behavior, number of nose pokes in the sucrose port were recorded during CS presentation; nose pokes and licks were also registered during the ITI period.

### Unexpected reward omission and extinction sessions

In an additional session, animals were subjected to an unexpected reward omission session, in which reward was omitted in 25% of the trials (8/30 trials, randomly assigned). This session was followed by 3 extinction sessions, one session per day, in which CS1 was presented but no sucrose was given.

### Aversive Pavlovian conditioning

After habituation to the chamber, mice started a 3-day aversive Pavlovian conditioning protocol. All sessions started with 60 seconds of habituation period, with the house light on. The CS2 consisted of an 80dB, 2kHz tone plus a cue light, paired with a mild foot shock (0.5mA, over 1 second) (US2). During the first conditioning session, mice were exposed to 10 CS-US pairings. Each trial consisted of a random ITI (35-50 seconds) followed by a 10 second tone, which was immediately followed by electric foot shock delivered through the stainless-steel grid floor. On the second day (US omission), mice performed 30 trials in which 10 out of 30 tones (randomly assigned) were not followed by an electric foot shock. On the third day, animals received a US extinction session, in which they received 30 trials with only CS exposure, but without US delivery (protocol adapted from^57^). Part of the animals received 7 additional days of US extinction. All sessions were recorded with webcams. To assess CS-triggered conditioned responses, two researchers evaluated freezing behavior during the CS period from all sessions in a blind manner. The freezing response was defined as the time (seconds) that mice spent immobile (lack of any movement including sniffing) except respiration during the CS period and calculated as percentage of total cue time ((freezing time x 100)/cue duration). Researchers that performed freezing analysis were blind to group and condition.

### Unexpected shock omission and extinction sessions

In an additional session, animals were subjected to an unexpected shock omission session, in which shock was omitted in 25% of the trials (4/12 trials, randomly assigned). This session was followed by 9 days of extinction, in which CS2 was presented but no shock was delivered.

### Optogenetic manipulation

For all optogenetic experiments using ChR2 for optical excitation, 5mW of blue light (at the tip of the fiberoptic) was generated by 473nm DPSS laser (CNI Laser, Changchun, China) and unilaterally delivered to mice through fiberoptic patch cords (0.22NA, 200μm diameter; Thorlabs, Newton, NJ, USA) that were attached to the implanted ferrule. For optogenetic experiments using eNpHR for optical inhibition, 5mW of yellow light (at the tip of the fiberoptic) was generated by 589nm DPSS laser (CNI Laser, Changchun, China) and unilaterally delivered as above. Laser output was controlled using a pulse generator (Master-8; AMPI, New Ulm, MN, USA) to deliver light.

### Aversive Pavlovian conditioning with optogenetic manipulation

#### Modulation during the conditioning session

The same protocol as the one described above was performed, with optical inhibition (10s of constant light 5mW at the tip of the fiber) being paired with CS2.

#### Modulation during the extinction phase

Mice were exposed to a conditioning session like the one described above. On the extinction sessions optical manipulation (excitation: 25ms light pulses of 20Hz for 10s; optical inhibition: 10s of constant light 5mW at the tip of the fiber) was paired with CS2 presentation, with no foot shock being delivered. Mice were exposed to 9 identical extinction sessions with optical manipulation.

### Locomotor activity with optogenetic manipulation

Locomotor activity was evaluated in an open field arena (43.2 cm x 43.2 cm) with transparent acrylic walls and white floor (Med Associates Inc., St. Albans, VT, USA). Briefly, mice were attached to an optical fiber connected to a laser (473 nm or 589 nm) and immediately placed in the center of the arena. Locomotion was monitored online over a period of 10 minutes (stimulation was given similarly as in the aversive Pavlovian conditioning: 10s of light stimulation followed by a 50s no stimulation interval). Distance traveled during the 10-minute session was used as indicator of locomotor activity. Average of all stimulation (0-10s) and post-stimulation (10-60s) periods is presented.

### Calcium imaging acquisition

GCaMP6f fluorescence signals were acquired using a miniature integrated fluorescence microscope system (Inscopix, Palo Alto, CA) through GRIN lenses implanted in the NAc on freely behaving mice. Before each imaging session, the miniaturized microscope was attached to the baseplate, by gently restraining the mouse. The analog gain (3.2-6) and LED output power (0.8-1.5 mW) of the microscope were set to be constant for the same subject across imaging sessions. The microscope focus was adjusted such that the best dynamic fluorescence signals were at the focal plane, which was subsequently kept constant across imaging sessions. To synchronize behavioral events with imaging acquisition, the Data Acquisition Box of the nVoke Imaging System (Inscopix, Palo Alto, CA) was triggered by the pyControl behavioral software. Compressed gray scale images were then recorded at 20 frames per second and with spatial down sampling by a factor of 4. Timestamp of each video frame was synchronized with and recorded by the pyControl behavioral acquisition system. Calcium imaging videos were acquired during the sucrose and shock sessions, during every day of the appetitive Pavlovian conditioning for half of the animals, while the remaining animals were recorded on days 1, 2, 3, 4, 5, 6, 7 and 10. For the aversive Pavlovian conditioning, we registered all days.

### Calcium imaging data processing

#### Data preprocessing

Using the Inscopix Data Processing Software (IDPS), we performed a field of view cropping to remove marginal areas and fixed this region for all task recordings for each animal. Subsequently, using IDPS, we applied a low-pass filter for noise reduction and a first motion correction using standard IDPS settings, exporting the result as a single TIFF image stack. Next, we applied a second motion correction using the CaImAn toolbox in Python58, which incorporates the NoRMCorre algorithm59. This algorithm performs a fast non-rigid motion correction that collectively optimizes artifact removal caused by movements. Following these corrections, we used the extended constrained non-negative matrix factorization developed for one-photon analysis (CNMF-E)60 in CaImAn to perform source separation (automatic identification of regions of interest (ROIs)) and obtain denoised and deconvolved fluorescence temporal activity, termed F. This, combined with a custom Python script that estimates the noise level of temporal traces (F_0) through the CaImAn internal function GetSn, yielded the normalized signals, termed ΔF/F_0. Subsequently, we manually inspected all automatically detected ROIs to eliminate irregular shapes or noisy calcium activity and to confirm that the identified ROIs corresponded to cells. Examples of ROIs and normalized signals obtained from a field of view, after cropping to remove marginal areas, are shown in Figure 1D.

#### Cell registration

After manual inspection, we extracted the spatial footprints of confirmed cells from CNMF-E for each session. Given a set of chronologically ordered sessions, we employed two methods for cell tracking: 1) CellReg^61^: a MATLAB script package that aligns spatial footprints across sessions using translation and rotation methods, with the first session as the reference map. For each cell pair, CellReg calculates a probability (P_same) of being the same cell, based on spatial correlations and centroid distance. Using default CellReg settings, a cell was considered tracked if P_same > 0.5; 2) Internal methods of the CaImAn toolbox: the *register_multisession* function uses spatial footprints of each cell and spatial correlation maps of each session to obtain an intersection over union metric for calculating distances between different cells in different sessions. It then solves a linear assignment problem using the Hungarian algorithm to determine the most likely cell pairings across sessions. Like CellReg, *register_multisession* uses the first session as the reference map. We adjusted the input parameters, setting the following variables “maximum distance considered” to 0.9 and the “max distance between centroids” to 100, providing greater flexibility compared to CellReg. We combined the results from both tracking methods and confirmed them through manual inspection.

### Calcium Data analysis

#### Alignment of activity to behavioral events

##### Sucrose and condensed milk

The occurrence of sucrose or condensed milk consumption events was defined as the detection of the first poke followed by licks (1 licking episode was defined as having at least 2 lick events occurring less than 250 ms apart) in the behavior box within 10 seconds of sucrose delivery. This criterion ensured that events were accurately reflecting consumption. Since we observed a high correlation of the first poke with licking behavior, indicative of consumption, in the Pavlovian conditioning, activity data is aligned to the first poke after sucrose delivery. For each event, a 3-second window pre-event and a 3-second window post-event were analyzed.

##### Foot shock exposure

For foot shock event, a 3-second window pre-foot shock and a 3-second window post-foot shock were analyzed.

##### Tail lift

Tail lifts were similarly marked with an external TTL signal triggered manually. For each tail lift, a 3-second window pre-TTL and a 3-second window post-TTL were analysed.

### Permutation test

To classify the response of a cell to a stimulus, we analyzed changes in their average signal ΔF/F_0. We used a permutation test where the fluorescence signal was shuffled across the 6-second window (3 seconds before and after stimulus presentation) 1000 times. If the absolute difference between the average pre- and post-stimulus signal in a real trial was statistically less 5% than observed in the 1000 shuffled data, the neuron was considered excited when the post-stimulus signal increased and inhibited when it decreased. Neurons with non-significant differences were classified as non-responsive.

### Signal normalization (z-score)

For subsequent analyses, we computed the z-score to represent the intensity of a cell’s response to a stimulus (CS or US). The z-score is calculated as z(t) = (F(t) -F_avg) / F_SD, where F(t) represents the normalized fluorescence signal ΔF/F_0, and F_Avg and F_SD are the mean and standard deviation of ΔF/F_0 measured across all 3-second pre-stimulus baselines in all trials, respectively. For trial-by-trial analyses, the z-score is computed using F_avg and F_SD measured only for the evaluated trial and in the 6 seconds immediately preceding the stimulus. In cases where the baseline contains only spurious activity, such as decay transients of the calcium signal or null activity, F_SD is set to the standard deviation over all 6-second pre-stimulus baselines, which serves as the expected natural standard deviation.

### Heatmaps

Heatmaps were constructed using the average z-score of individual cell activity using the seaborn package in Python (seaborn.heatmap). The interval was set from −3 to 3 seconds, where zero indicates the stimulus onset (CS or US). When heatmaps were shown for the activity of all cells recorded in each session for two stimuli, the responses were sorted in descending order based on the magnitude z-score of the first stimulus, and the same cell order was maintained for the heatmap of the second stimulus. When heatmaps were displayed for tracked cells, even for different sessions, the responses were aligned in descending order to the first stimulus of the first reference session. In all cases, the color bar was configured with ‘extend’ in seaborn.heatmap, indicating that the color scale extends beyond the upper and lower bounds of the data. The color scale was set to have an upper bound of 1.5 and a lower bound of −0.5 for the z-score.

### PSTHs and AUCs

After classifying the cells responses to a stimulus, we constructed peristimulus time histograms (PSTHs) for each group of response types (excited, inhibited, and non-responsive). For each group, individual z-scores were averaged, and the mean with standard error of the mean (SEM) was plotted. Additionally, the area under the curve (AUC) was calculated for each group during the 3 seconds following stimulus onset using the Python package Scikit-learn (function sklearn.metrics.auc). These methods were also applied to obtain PSTHs and AUCs for each cluster of cells based on their responses on day 1 of appetitive and aversive conditioning.

### Percentage of persisted responses

After classifying a cell’s response to a stimulus as excited, inhibited, or non-responsive based on the mean across trials in the permutation test, we calculated the percentage of persisted responses. This involved determining the fraction of trials in which the same response was obtained when the permutation test was applied to each single trial. We defined that >= 70% of persisted responses indicates that the cell’s response to the stimulus is consistent (Extended Data Fig. 2E-F).

### Shannon’s entropy

To assess the variability of neuronal responses to each stimulus, we used Shannon’s entropy. For each presentation of the same stimulus, the resulting neuronal responses formed a series, which was subsequently utilized to calculate Shannon’s entropy using the expression -Σ_i_p(i)log2[p(i)], where p(i) denotes the probability of observing a specific type of response (e.g., excitatory, inhibitory, or non-responsive). In the most uncertain (random) scenario, where the neuron could produce any response for each stimulus presentation, p(i) = 1/3, resulting in an approximate maximum entropy of 1.58. Conversely, in the deterministic scenario, where the neuron consistently responds in the same manner for every stimulus presentation, the entropy is zero. In the plots, the dashed lines indicate entropy values corresponding to 33, 50, 60, 70, 80, 90 and 100 % of repeated responses, with the remaining being distinct and equiprobable.

### Single cell decoder analysis

We used the average z-score of the cell’s 3-second response to each stimulus presentation for single-cell decoding analyses. Given a cell with M_A trials responding to stimulus A and M_B trials responding to stimulus B, we employed balanced datasets for training a linear SVM model (using sklearn.svm.LinearSVC in Python with C = 0.8). If M_A is greater than M_B, we randomly subsampled M_B trials from M_A. Otherwise, we randomly subsampled M_A trials from M_B. From the actual data, we used 75% of the trials for each stimulus for training and reserved the remaining 25% for machine accuracy testing. To generate the corresponding shuffled data, we conducted 100 permutations of the ΔF/F_0 signal for each window from −6 to 3 seconds relative to stimulus onset. We then proceeded with the z-score calculation for trial-by-trial as described previously. This procedure was mirrored to create a classifier using the actual data to decode the shuffled data. Finally, we generated 1000 independent machines for both the actual test data and the shuffled data. The average of the accuracy test results on these machines represents the single cell decoder accuracy. In Fig. 3F-G, the decoding accuracy of each neuron (neuron ID on X axis) on day 1 was plotted following the color code of the accuracy (red-high, blue – low), and gray dots represent the decoding accuracy of those same neurons on day 5 (left) or day 10 (right). For Fig. 3H-I, neurons were divided into 20% best decoders, or 20% worst decoders based on day 1 activity.

### Population decoder analysis

For population-level decoding analyses, we use a population vector of size N, where N is the total number of cells responding to different stimuli. Each element in this vector represents the average z-score of a cell’s 3-second response to a stimulus presentation. Each trial generates a unique population vector that serves as a representation of the presented stimulus. A stimulus presentation trial generates a population vector that represents that stimulus. Like the single-cell analysis, we ensured balanced datasets for the SVM model by selecting an equal number of population vectors from each stimulus category. As before, we selected an equal number of population vectors from each stimulus to generate datasets for the SVM model. This way, we allocated 75% of the population vectors for model training and the remaining 25% for machine accuracy testing. We generated 1000 independent machines for both the actual test data and the shuffled data. The average of the accuracy test results on these machines represents the population decoder accuracy. For the special case in which the SVM model was trained for the CS1, US1, CS2, and US2 stimuli (Figure 6), we used the sklearn.svm.SVC function in Python with the following arguments: C=0.8, kernel=’linear’, and decision_function_shape=’ovo’ (one-vs-one).

### Correlation analysis

To assess individual variability in response dynamics to the same stimulus within a single session, we constructed an M x M matrix for each neuron, where M is the number of trials for the same stimulus. Each element Mij of the matrix was computed as the Pearson correlation coefficient between the 3-second z-scored responses to the stimulus for trial i and trial j. The single neuron activity correlation was then derived as the mean of the elements located above the main diagonal of matrix M, representing the average Pearson correlation coefficient across all combinations of trial pairs. This is exemplified in Figure 1L-M, which shows the distribution of this correlation for neurons for US1 and US2.

In addition to single neuron activity correlation, we also assessed the tuning curve correlation, which quantifies the similarity in response profiles of individual neurons to the same stimulus across multiple presentations within a session^61^. For each trial i, we constructed an N x T matrix of z-scores, where N is the number of neurons recorded from an animal and T is the number of time frames in the analysis period (3 seconds post-stimulus, acquired at 20 Hz, resulting in T = 60). The tuning curve correlation between trials i and j was then calculated as the median of the Pearson correlation coefficients between corresponding neurons across trials. Extended Data Fig. 2I illustrates how the tuning curve correlation changes on average as we compare trials with increasing separation (“distance between trials”). For this figure, especially for sucrose consumption (US1), we only showed 13 consumptions per animal. This is because consumption varied between animals, and 13 represented the least common number of consumptions observed.

To evaluate the correlation in individual response dynamics to the same stimulus and across different sessions, for each tracked neuron, we calculated the Pearson correlation coefficient between the average signal of z-scored responses (3-second window) to the stimulus across distinct sessions. This assesses how consistently individual neurons respond to the same stimulus over time. For instance, in Extended Data Fig. 4I-J, the distributions of these correlations of all tracked neurons for different pairs of days during appetitive conditioning are shown, with the averages of the distributions of each pair being highlighted by vertical lines.

Furthermore, we examined correlation between the CS-US stimuli at the population level. For each neuron, we averaged its responses across all trials and within a 3-second window for each stimulus. For example, this data is visualized in Extended Data Fig. 3D-E, where each data point represents an average response pair CS-US. To quantify the association between these paired responses, we calculated the Pearson correlation coefficient and performed linear regression to identify possible trends in the data. We also measured the angle between the closest points of each response pair. This metric, in the form of a histogram, allows from another perspective to compare the evolution of these data when analyzing different sessions. These analyses were not only performed on all neurons but also on a subset of “CS-US responsive” neurons identified through a permutation test. This approach allowed us to concentrate on neurons that showed significant activity changes in response to both stimuli.

We also evaluated the population vector correlation (PV correlation), which is a measure of similarity between multiple responses to the same stimulus across a neuronal population. This measure has been applied in detecting drift in overall activity patterns over time, comparing “within-session” (single session) and “between-sessions” (different sessions) correlations (for more details, see^61^). Briefly, for each stimulus (CS or US) and tracked neurons, we split each session into two blocks (first and second half of trials). For each neuron, we then averaged the z-scores across all frames within each block. In these blocks, a population vector is a set of average z-score values at a given frame, thus, a block is formed by an N x T matrix, where N is the number of neurons and with T = 60 frames (3 seconds). We then calculate the PV correlation between pairs of blocks by averaging the Pearson correlation coefficients between their respective population vectors at each frame. When the two blocks are from the same session, the values of PV correlation are added to the “within-session” group, and when the blocks are from different sessions, the values of PV correlation are added to the “between-sessions” group. Finally, Fig. 3M-N displays the average values of these groups across animals for D1- and A2A-neurons.

### Cluster analysis and representation

For cluster analysis, we concatenated the z-scores of the average activity over all trials from −3 to 3 seconds (where zero corresponds to stimulus onset) of both the CS and its corresponding US, totaling 12 seconds. Treating each timepoint (frame) as an independent dimension, we applied Principal Component Analysis (PCA, *sklearn.decomposition.PCA* function in Python) to a matrix of dimensions N x D, where N is the number of cells and D is the total number of dimensions. In our case, this resulted in 240 dimensions (12 seconds x 20 Hz, the sampling rate). Subsequently, we performed dimensionality reduction, selecting enough Principal Components (PCs) to account for at least 80% of the observed variance in our data (Extended Data Fig. 6A). This typically ranged from 12 to 15 PCs, which were then subjected to the K-means clustering method using the *sklearn.cluster.KMeans* function in Python. To determine the optimal number of clusters (n_c) for the reduced data, we varied n_c from 2 to 10 and performed the following assessments: 1) Identified the n_c that marked the onset of stability in the average cosine similarity computed between cluster elements and the average cluster activity (Extended Data Fig. 6B); 2) Identified the n_c that yielded a peak in silhouette score analysis using the Euclidean metric (Extended Data Fig. 6C). The concordance of these two measures defined the n_c to be employed in subsequent analyses. As an illustrative example, we examined the average z-scores of D1 neurons recorded on day 1 of appetitive conditioning, and this analysis revealed that n_c = 3 was the optimal number of clusters for this dataset. To visualize the identified clusters, in Extended Data Fig. 6D, we utilized two dimensionality reduction techniques: PCA (a linear method) and t-Distributed Stochastic Neighbor Embedding (t-SNE, a non-linear method), and for both, we use the first two components, and each color represents a cluster.

### Neuronal trajectory

To investigate the evolution of neural activity patterns following positive and negative valence stimuli, we conducted neural trajectory analysis. This approach tracks changes in a tracked population response over time. For each tracked cell, we concatenated z-scores of average activity across all trials within a 3-second window after stimulus onset. To analyze US1 and US2 in Fig. 1R-S, we generated a matrix of size N x 120 (2 stimuli, each comprising 60 frames). Similarly, to analyze CS1, CS2, US1, and US2 in Fig. 6G-I, we generated a matrix of size N x 240. Each matrix was generated per animal, where N represents the number of cells. To visualize the overall activity dynamics, we reduced the matrix dimensionality to two principal components (PC) using PCA treating each cell as an independent dimension. We then smoothed these components using one-dimensional Gaussian convolution, resulting in neural trajectories shown as solid lines in these figures.

To quantify the dissimilarity between trajectories over time for different stimuli, we calculated the Euclidean distance between their corresponding PC values at each time point. Specifically, we computed distances between (PC1_EST1, PC2_EST1) and (PC1_EST2, PC2_EST2) at each time point, where EST corresponds to CS or US.

Furthermore, we visualized single-trial variability by plotting individual trial trajectories as dashed lines (Fig. 1R-S and Fig. 6G-I). These trajectories were obtained by applying the same PCA transformation matrix used for average activity data. Notably, we only plotted 7 trials per stimulus (the maximum number of trials for footshock) to maintain a balanced representation, similar to the population decoder analysis.

### Optogenetic manipulation

For all optogenetic experiments using ChR2, 5mW of blue light (at the tip of the fiberoptic) was generated by 473nm DPSS laser (CNI Laser, Changchun, China) and unilaterally delivered to mice through fiberoptic patch cords (0.22NA, 200μm diameter; Thorlabs, Newton, NJ, USA) that were attached to the implanted ferrule. For optogenetic experiments using eNpHR, 5mW of yellow light (at the tip of the fiberoptic) was generated by 589nm DPSS laser (CNI Laser, Changchun, China) and unilaterally delivered. Laser output was controlled using a pulse generator (Master-8; AMPI, New Ulm, MN, USA) to deliver light.

Optogenetic manipulation of D1- or D2-MSNs was time-locked to cue onset on each trial and lasted the entire period of the cue (10 seconds). Stimulation was performed during the conditioning day (Extended data Fig. 10) or during the nine days of extinction (Fig. 7).

### Sacrifice and brain sectioning for histological analysis

At the end of all behavior procedures, mice were deeply anesthetized by a mixture of ketamine/medetomidine. Animals were then transcardially perfused with 0.9% saline, followed by 4% paraformaldehyde (PFA) solution. After, whole heads with the lenses or the optic fibers attached were immersed for 48h in 4% PFA so that the lens track is clearly visible for histological analysis. Next, brains were extracted and then rinsed and stored in 30% sucrose at 4°C until sectioning.

Sectioning was performed coronally, in 40μm slices, on a vibrating microtome (VT1000S, Leica, Germany) and slices were stored at 4°C on 12-well plates (or long-term storage in cryoprotectant solution at −20°C) until use. Slices from the area of interest (NAc) were selected using the Mouse Brain Atlas^62^.

### Immunohistochemistry

To assess GCaMPf6 expression in D1-cre and A2A-cre mice, brain slices containing NAc sections were washed with phosphate buffered saline (PBS), and then permeabilizated with PBS-Triton 0.3% (PBS-T 0.3%). Blocking was performed for 1h using 5% Fetal Bovine Serum (FBS; Invitrogen, MA, USA) in PBS-T at RT. The primary antibody goat anti-GFP (1:750; ab6673, ABCAM, Cambridge, UK), was incubated overnight at 4°C with agitation, followed by PBS-T washes and subsequent incubation with the secondary fluorescent antibody Alexa Fluor® 488 donkey anti-goat (1:500; A11055, Invitrogen, Carlsbad, CA, USA). All antibodies were diluted in PBS-T with 2% FBS. Slices were washed with PBS-T, incubated with 4’,6-Diamidino-2-Phenylindole Dihydrochloride (DAPI, 1:1000; 62248, Thermo ScientificTM, Waltham, MA, USA) for nucleus staining, washed with PBS and mounted using Permafluor (Invitrogen, MA, USA). Slides were stored at 4°C and kept protected from light.

### Image acquisition and analysis

Images from the NAc of D1-cre and A2A-cre mice were collected in an inverted confocal microscope (Olympus FV3000, Tokyo, Japan). About 6 slices per animals were used for each analysis. Optic fiber placement was assessed to confirm if the activity detected was from the NAc region. For that, slices where the optic fiber was detected were classified according to the Mouse brain atlas^62^ to estimate the stereotaxic coordinates.

### Statistical analysis

Part of the statistical analysis was performed in GraphPad Prism 9.0 (GraphPad Software, Inc., La Jolla, CA, USA). Prior to any statistical comparison between groups, a Tukey test was performed to assess presence of outliers. Normality was assessed in all data analyzed by using the Kolmogorov-Smirnov (KS) test. Parametric tests were used whenever KS test > 0.05. If normality assumptions were not met, non-parametric analysis (Mann-Whitney or Wilcoxon test) was performed.

For behavior, two-way analysis of variance (ANOVA) for repeated measures was used to assess learning in D1- and A2A-cre mice (factors used: nose pokes during CS *versus* nose pokes during ITI, across days of training); and to analyze percentage (%) of freezing throughout the trials on day 1. Bonferroni’s *post hoc* multiple comparison test was used for group differences determination. One Way ANOVA was done to compare the poke probability at the ITI or CS period on days 1, 5 and 10 of learning in the appetitive Pavlovian conditioning task. One Way ANOVA was also done to compare the latency to compare the time between delivery of the reward and the first poke on days 1, 5 and 10 of training. Statistical analysis between two time-points was made using two-tailed paired Student’s t-test, to compare percentage of freezing in the first and last trials of the aversive conditioning. Two-way ANOVA for repeated measures was performed to compare the percentage of freezing across days of shock extinction.

Data are presented as mean ± standard error of mean (SEM). Statistical significance was considered for p ≤ 0.05. All statistical data of the main figures are depicted in Supplementary Table 1. All statistical data of the extended data figures are depicted in Supplementary Table 2.

### Data and code availability

The preprocessed data utilized in this study, as outlined in the Methods section, are openly accessible and deposited on the Zenodo platform: [link provided after acceptance of the manuscript]. Workflow and toolboxes developed by authors for calcium imaging data analysis will be openly available on GitHub repository [link provided after acceptance of the manuscript].

## Acknowledgments

This work was funded by the European Research Council (ERC) under the European Union’s Horizon 2020 research and innovation programme (grant agreement No 101003187) and by the “la Caixa” Foundation (ID 100010434), under the agreement LCF/PR/HR20/52400020. The work was also funded by a Bial Foundation grant (175/2020). Part of the work received funding from the FCT under the scope of the projects PTDC/MED-NEU/4804/2020 (DOI: 10.54499/PTDC/MED-NEU/4804/2020), PTDC/SAU-TOX/6802/2020 (DOI: 10.54499/PTDC/SAU-TOX/6802/2020), 2022.02201.PTDC (DOI: 10.54499/2022.02201.PTDC) and 2022.01467.PTDC (DOI: 10.54499/2022.01467.PTDC).

CS-C, BC and LP have Scientific Employment Stimulus contracts from the Portuguese Foundation for Science and Technology (FCT) (CEECIND/03887/2017 (DOI:10.54499/CEECIND/03887/2017/CP1458/CT0027); CEECIND/03898/2020 (DOI:10.54499/2020.03898.CEECIND/CP1600/CT0015); CEECINST/00077/2018 (DOI:10.54499/CEECINST/00077/2018/CP1640/CT0003)). AVD and RC have FCT PhD grants (SFRH/BD/147066/2019; 2022.12973.BD).

This work was also supported by a FEBS (Federation of European Biochemical Societies) Excellence Award, IBRO Early Career Award and a “Maria de Sousa” Award attributed to Carina Soares-Cunha.

Host laboratory is funded by National funds, through FCT -project UIDB/50026/2020 (DOI 10.54499/UIDB/50026/2020), UIDP/50026/2020 (DOI 10.54499/UIDP/50026/2020) and LA/P/0050/2020 (DOI 10.54499/LA/P/0050/2020).

## Contributions

CS-C and AJR conceived the project, designed the experiments and supervised the research; AVD performed the majority of the experiments; TTAT analyzed most of the data; GJM supported in the collection of 1-photon images; RC performed histological confirmation; BC supported in optogenetic experiments; RG performed analysis of aversive Pavlovian conditioning data; RG was responsible for colony management and generation of transgenic mice; MW helped in the optogenetic experiment data analysis; LP provided support to imaging data collection and analysis; RMC and NS supervised the research.

## Ethics declaration

The authors declare no competing interests.

